# Reduced cell-substrate adhesion formation promotes cell migration in *Dictyostelium*

**DOI:** 10.1101/2024.03.19.585764

**Authors:** Julio C. Fierro Morales, Thu Nguyen, Sabin Nepal, Chandler Redfearn, Bruce K. Gale, Margaret A. Titus, Minna Roh-Johnson

## Abstract

Many cells adhere to the extracellular matrix for efficient cell migration. This adhesion is mediated by focal adhesions, a protein complex linking the extracellular matrix to the intracellular cytoskeleton. Focal adhesions have been studied extensively in Metazoan mesenchymal cells, but recent research in physiological contexts and amoeboid cells suggest that focal adhesion regulation differs from the mesenchymal focal adhesion paradigm. While focal adhesion machinery predates the origin of Metazoans, focal adhesion formation and regulation during non-Metazoan cell migration is largely unexplored. We used *Dictyostelium discoideum* to investigate novel mechanisms and the evolution of focal adhesion regulation, as *Dictyostelium* are non-Metazoans that form cell-substrate adhesion structures for migration. We show that PaxillinB, the *Dictyostelium* homologue of Paxillin, localizes to dynamic cell-substrate adhesions. As expected, PaxillinB mutations decreased the number of cell-substrate adhesions. Unexpectedly, however, decreased cell-substrate adhesion number led to an increase in cell migration speed. These findings are in direct contrast to Paxillin function at focal adhesions and regulation of cell migration in mammalian cells, challenging the established focal adhesion model and providing insight into the evolution of cell-substrate adhesions and Paxillin function during cell migration.

**Significance Statement:**

- Focal adhesions are understudied in non-mesenchymal, non-Metazoan systems.
- The authors characterize PaxillinB, the *Dictyostelium* homologue of Paxillin, during *Dictyostelium* cell migration. Reducing cell-substrate adhesion number via PaxillinB mutations lead to decreased cell adhesion and, surprisingly, increased cell migration speeds.
- These findings suggest *Dictyostelium* cells form cell-substrate adhesions that act as molecular “brakes” during cell migration and provides unique insight into the evolution of Paxillin as a regulator of cell-substrate adhesion function and formation during cell migration.

## INTRODUCTION

Single-cell crawling migration is a pivotal aspect of several developmental, immunological, and disease processes, with cells being able to switch between distinct forms of motility depending on the environmental context (De Pascalis and Etienne-Manneville, 2017; Merino-Casallo et al., 2022; Petrie and Yamada, 2016; Trepat et al., 2012). This ability to switch between different forms of crawling motility is important for cells to adapt to unique, rapidly changing environments and is conserved across the species tree of life (Brunet et al., 2021; De Pascalis and Etienne-Manneville, 2017; Fritz-Laylin and Titus, 2023; Pawluchin and Galic, 2022; Yamada and Sixt, 2019). Crawling cell migration is often categorized into two overarching forms: amoeboid migration and mesenchymal migration, although cell migration should be considered as more of a spectrum of states (Kunwar et al., 2006; Te Boekhorst et al., 2016; Yamada and Sixt, 2019). Amoeboid motility, which is typically associated with non-Metazoan amoebae and immune cells such as neutrophils and T cells, is defined by low-to-no adhesion to the surrounding extracellular matrix (ECM), formation of both actin-dependent and independent (blebbing) protrusions, and increased actomyosin contractility (Friedl and Wolf, 2010; Lämmermann and Sixt, 2009; Pawluchin and Galic, 2022; Rodriguez-Hernandez et al., 2016; Schick and Raz, 2022; Yamada and Sixt, 2019). In contrast, mesenchymal migration is characterized by strong adhesion to the ECM, a rich actin network at the leading edge, and ECM proteolysis in mammalian cells such as fibroblasts, neural crest cells, and several cancer cell lines (Kunwar et al., 2006; Merino-Casallo et al., 2022; Pawluchin and Galic, 2022; Yamada and Sixt, 2019).

Research focused on mesenchymal migration has shown that cells use adhesion structures called focal adhesions. Focal adhesions are multilayered nanostructures composed of hundreds of proteins that physically link the surrounding ECM to the intracellular actin cytoskeleton (Case and Waterman, 2015; Kanchanawong et al., 2010; Lo, 2006). Integrin heterodimers are transmembrane proteins that physically link focal adhesions to the ECM. Upon integrin interaction with the ECM, a series of mechanosensitive and biochemical signals regulate cycles of focal adhesion assembly and disassembly. This cycle, in turn, promotes efficient mesenchymal cell migration (referred to as adhesion-based migration going forward) (Geiger et al., 2009; Lock et al., 2008; Oakes and Gardel, 2014). There is extensive literature dissecting focal adhesions during adhesion-based migration, and the majority has focused on mesenchymal cells in flat, *in vitro* 2D systems (Fierro Morales et al., 2022). Additionally, nearly all focal adhesion research has been conducted in Metazoan models – as focal-adhesion based migration has classically been considered unique to Metazoans – while amoeboid motility is believed to be ubiquitous across Eukaryotes (Brunet et al., 2021; Devreotes and Zigmond, 1988; Fritz-Laylin et al., 2017; Fritz-Laylin and Titus, 2023; Velle and Fritz-Laylin, 2019; Velle and Fritz-Laylin, 2020). Studies in these *in vitro,* 2D Metazoan systems have led to an established model of focal adhesion components, assembly, and dynamics characterized by large, mature focal adhesion structures composed of core components that enable the generation of traction forces and a slower rate of migration compared to other forms of motility such as amoeboid migration (Case and Waterman, 2015; Gardel et al., 2010; Kanchanawong et al., 2010; Liu et al., 2015; Pankova et al., 2010; Parsons et al., 2010).

Recent research dissecting focal adhesion formation and regulation in other contexts and cell types, however, has led to questions as to the criteria defining focal adhesions and the cells and organisms that are capable of forming them for cell migration. Work in under-investigated contexts such as *in vivo* models of mesenchymal cells, physiologically-mimicking environments, and immune cells has characterized focal adhesion structures that differ from the canonical model in terms of dynamics, composition, and regulation (Caillier et al., 2024; Xue et al., 2023; Zhang et al., 2023). These results suggest that under-studied systems are ripe for discovering novel aspects of focal adhesion biology that deviate from the established paradigm. One context for investigating under-studied forms of focal adhesion regulation and formation during cell migration is in non-Metazoan species. Focal adhesion machinery was previously believed to be a Metazoan-specific innovation that arose in Urmetazoa, the last common ancestor of Metazoa, to help enable the multicellular transition for the origin of animals (Abercrombie et al., 1971; Hynes and Zhao, 2000; King, 2004; Sebé-Pedrós et al., 2010). Genomic analyses of non-Metazoan organisms following the advent of next-generation sequencing technology, however, identified putative homologues of various core components of focal adhesions in organisms as evolutionarily distant as Amoebozoa, suggesting that this machinery originated prior to the divergence of Metazoa (Fierro Morales et al., 2022; Hynes and Zhao, 2000; Rokas, 2008; Sebé-Pedrós et al., 2010). Intriguingly, these analyses suggest that not all components of the canonical focal adhesion toolkit evolved concurrently, with molecules originating at different points across the species tree of life (Fierro Morales et al., 2022; Sebé-Pedrós et al., 2010). Furthermore, focal adhesion protein homologues are present in non-Metazoan species that are believed to predominantly use focal adhesion-independent migration (i.e. amoeboid-like migration). These seemingly paradoxical findings raise questions on if and how non-Metazoan organisms use focal adhesion-like structures for cell migration and how the composition, function, and regulation of putative non-Metazoan focal adhesions have evolved across the species tree of life.

Among the non-Metazoan species that possess homologues of focal adhesion molecules is the model Amoebozoan, *Dictyostelium discoideum. Dictyostelium* is a soil-dwelling amoeba that transitions from a unicellular to a multicellular state via single cell migration and subsequent aggregation. *Dictyostelium* has historically been utilized to study single cell amoeboid migration and chemotaxis, serving as a powerful comparative model for other amoeboid cells typically associated with adhesion-independent migration such as leukocytes (Devreotes and Zigmond, 1988; Friedl et al., 2001; Fukui and Inoue, 1997; Paluch et al., 2016). Recent studies characterizing conserved focal adhesion molecules in *Dictyostelium*, however, provide initial evidence indicating that these amoebae are capable of forming actin-rich focal adhesion-like structures (referred to as cell-substrate adhesions from this point onwards). These cell-substrate adhesions contain homologues of several focal adhesion proteins (Berman et al., 2000; Bukharova et al., 2005; Cornillon et al., 2008; Cornillon et al., 2006; Cornillon et al., 2000; Duran et al., 2009; Fey et al., 2002; Froquet et al., 2012; Hibi et al., 2004; Nagasaki et al., 2009; Niewöhner et al., 1997; Patel et al., 2008; Plak et al., 2016; Pribic et al., 2011; Tsujioka et al., 2004; Tsujioka et al., 2008), and generate traction forces on the substratum during migration, reminiscent of Metazoan focal adhesions (Copos et al., 2017; Iwadate and Yumura, 2008a; Mijanovic and Weber, 2022; Uchida and Yumura, 2004). Interestingly, homology analyses suggest that *Dictyostelium* possess putative homologues of some, but not all, focal adhesion components (Fierro Morales et al., 2022; Sebé-Pedrós et al., 2010), indicating that *Dictyostelium* is able to use cell-substrate adhesion-based migration despite lacking core proteins thought to be critical for focal adhesion formation and function. The lack of putative homologues of core focal adhesion components makes *Dictyostelium* an ideal model for investigating the evolution of focal adhesion composition, function, and regulation for single cell migration. In addition to serving as an intriguing model for comparing non-Metazoan and canonical Metazoan focal adhesion structures from an evolutionary lens, insight gained from further characterizing cell-substrate adhesions structures in *Dictyostelium* will help elucidate how focal adhesions are assembled and regulated for efficient migration in conditions where key components are missing. This in turn can guide research into differential modes of focal adhesion regulation and assembly relevant to physiological environments (Caillier et al., 2024; Xue et al., 2023; Zhang et al., 2023).

We sought to understand how *Dictyostelium* forms and regulates cell-substrate adhesions during single cell migration. We focused on characterizing PaxillinB, the *Dictyostelium* homologue of the focal adhesion scaffolding molecule Paxillin. Initial work investigating PaxillinB in *Dictyostelium* suggests that PaxillinB localizes to cell-substrate adhesions and is involved in adhesion to the underlying substrate (Bukharova et al., 2005; Nagasaki et al., 2009; Patel et al., 2008; Pribic et al., 2011). However, how or whether PaxillinB regulates cell-substrate adhesion dynamics during *Dictyostelium* cell migration is unclear. We dissected the functions of PaxillinB at *Dictyostelium* cell-substrate adhesions and identified key conserved domains required for PaxillinB localization to adhesion structures. Unexpectedly, however, we observed that reduced PaxillinB recruitment to cell-substrate adhesions *increased Dictyostelium* cell migration speed. Further investigation showed that this increase in cell migration speed was not due to a switch to an adhesion-independent form of migration or increased rate of cell-substrate adhesion turnover. Instead, we found that the increased migration phenotype was due to reduced formation of cell-substrate adhesions. These results are in contrast to how Paxillin regulates adhesion-based migration in Metazoan systems, shedding light on novel forms of focal adhesion regulation alternate to the canonical, mesenchymal model. These findings also further our understanding of the evolution of focal adhesions and suggest an ancestral role for Paxillin-positive cell-substrate adhesions in promoting cell adhesion at the expense of cell migration speed, as opposed to promoting cell migration as has been largely established in mammalian models.

## RESULTS

### *Dictyostelium discoideum* possesses putative homologues of key focal adhesion components

To investigate mechanisms of cell-substrate adhesion formation, function, and regulation in *Dictyostelium*, we first sought to identify which core focal adhesion components (as defined by focal adhesion research in mammalian systems) are present in *Dictyostelium*. While work using BLASTp to dissect sequence conservation identified putative homologues of core focal adhesion components in *Dictyostelium* and other non-Metazoan organisms (Sebé-Pedrós et al., 2010), incorporating novel domain architecture-based and structural-based homology tools can provide higher sensitivity for homologue detection, particularly for distantly evolutionarily-related proteins (Illergard et al., 2009; Lee and Lee, 2009; Potter et al., 2018; van Kempen et al., 2024). Thus, we used a combination of bidirectional homology queries using BLASTp (McGinnis and Madden, 2004), PHMMR (Potter et al., 2018), and FoldSeek (van Kempen et al., 2024) to evaluate sequence, domain architecture, and predicted structural homology, respectively.

Consistent with previous studies (Sebé-Pedrós et al., 2010), our analyses suggest that homologues of some core focal adhesion molecules, particularly those involved in scaffolding and force transduction, originated in Eukaryota, just prior to the divergence of Amoebozoans (Figure 1). Homologues of other proteins necessary for mammalian focal adhesion formation – such as the tyrosine kinases and integrins – appear to have evolved separately (Figure 1). We also sought to identify specific non-Metazoan organisms that have been shown to form cell-substrate structures through functional analyses and/or imaging-based studies (Figure 1A). We found that these organisms, including *Dictyostelium*, possess homologues of some, but not all, core focal adhesion molecules (Figure 1A). Specifically, *Dictyostelium* lacks putative homologues of the key tyrosine kinases focal adhesion kinase (FAK) and c-SRC (Figures 1A and 2A), with other work further suggesting a lack of tyrosine kinases altogether (Goldberg et al., 2006). Furthermore, *Dictyostelium* possesses the Sib (Similar to Integrin Beta) molecules, which are putative beta-integrin-like proteins composed of a mix of conserved mammalian beta-integrin domains – such as cytosolic NPxY motifs for Talin binding and von Wildebrand A domains with metal ion dependent adhesion site (MIDAS) motifs associated with ECM binding – as well as adhesion domain repeats that have only been additionally identified in bacterial substrate adhesion proteins (Cornillon et al., 2008; Cornillon et al., 2006; Lally et al., 1999). This domain architecture suggests Sib proteins are somewhat of a “hybrid” adhesion molecule, similar to other integrin-like proteins that have been identified in non-Metazoan organisms (Kang et al., 2021). While Sib molecules are hypothesized to serve as beta-integrin-like adhesion receptors and have been shown to bind to *Dictyostelium* Talin via their NPxY motifs, to the best of our knowledge, no additional work has been done to confirm this function. These homology results, combined with previous work (Berman et al., 2000; Bukharova et al., 2005; Cornillon et al., 2008; Cornillon et al., 2006; Cornillon et al., 2000; Duran et al., 2009; Fey et al., 2002; Froquet et al., 2012; Hibi et al., 2004; Nagasaki et al., 2009; Niewöhner et al., 1997; Patel et al., 2008; Plak et al., 2016; Pribic et al., 2011; Tsujioka et al., 2004; Tsujioka et al., 2008), suggest that *Dictyostelium* is capable of forming cell-substrate adhesions without possessing all the components necessary for mammalian focal adhesion formation. We next determined the mechanism of cell-substrate adhesion formation in this amoeba.

**Figure 1:**
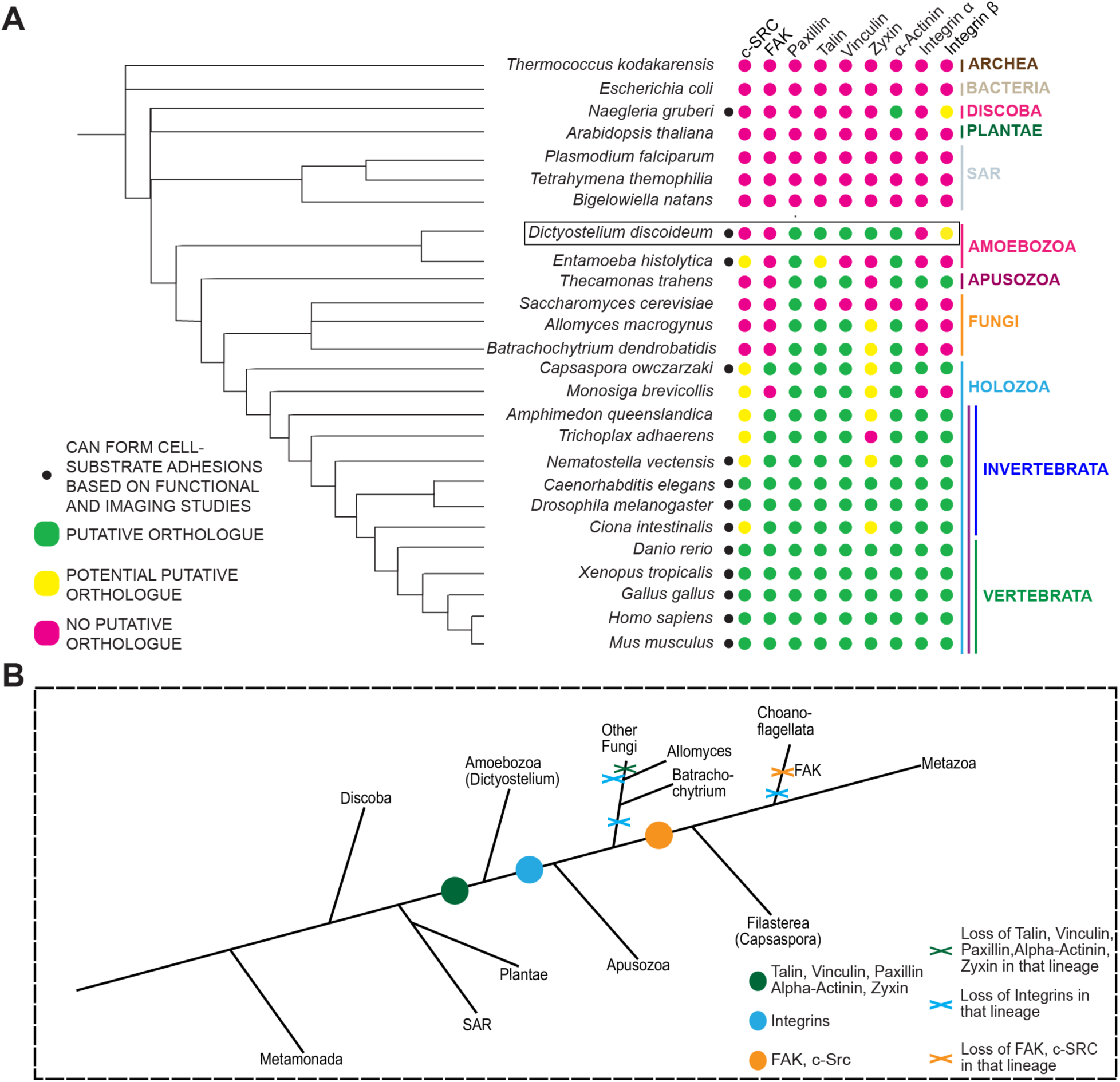
The model Amoebozoan *Dictyostelium discoideum* possesses putative homologues of some, but not all, core focal adhesion components. (A) A bidirectional homology pipeline using pHMMR, FoldSeq, and previously described BLASTp methods (Sebé-Pedrós et al., 2010) was utilized to map the presence and absence of homologues of core Metazoan focal adhesion components across evolutionary space. The presence and absence of these homologues is mapped against a species tree derived and expanded from previous work (Velle and Fritz-Laylin, 2019). Species with a black dot have been shown to form focal adhesions or focal adhesion-like structures either via imaging studies or functional analyses (Bukharova et al., 2005; Guler et al., 2021; Jahnel et al., 2014; Jamerson et al., 2012; Loveless et al., 2017; Parra-Acero et al., 2020; Santiago-Medina et al., 2012; Singiser and McCann, 2006; Stubb et al., 2019; Tavares et al., 2005; Turner et al., 1990; Xue et al., 2023; Zhao et al., 2022). Importantly, a lack of a black dot does not indicate that species is not capable of forming focal adhesions despite possessing core components; rather, it means that to the best of our knowledge, there is no published literature confirming focal adhesion or focal adhesion-like structure formation via imaging or functional analyses. (B) Schematic demonstrating the putative origin of core focal adhesion components across the species tree of life. Ours and other evolutionary analyses suggests some, but not all, core components of the focal adhesion machinery originated just prior to the divergence of Amoebozoa. Adapted from (Sebé-Pedrós et al., 2010).

**Figure 2:**
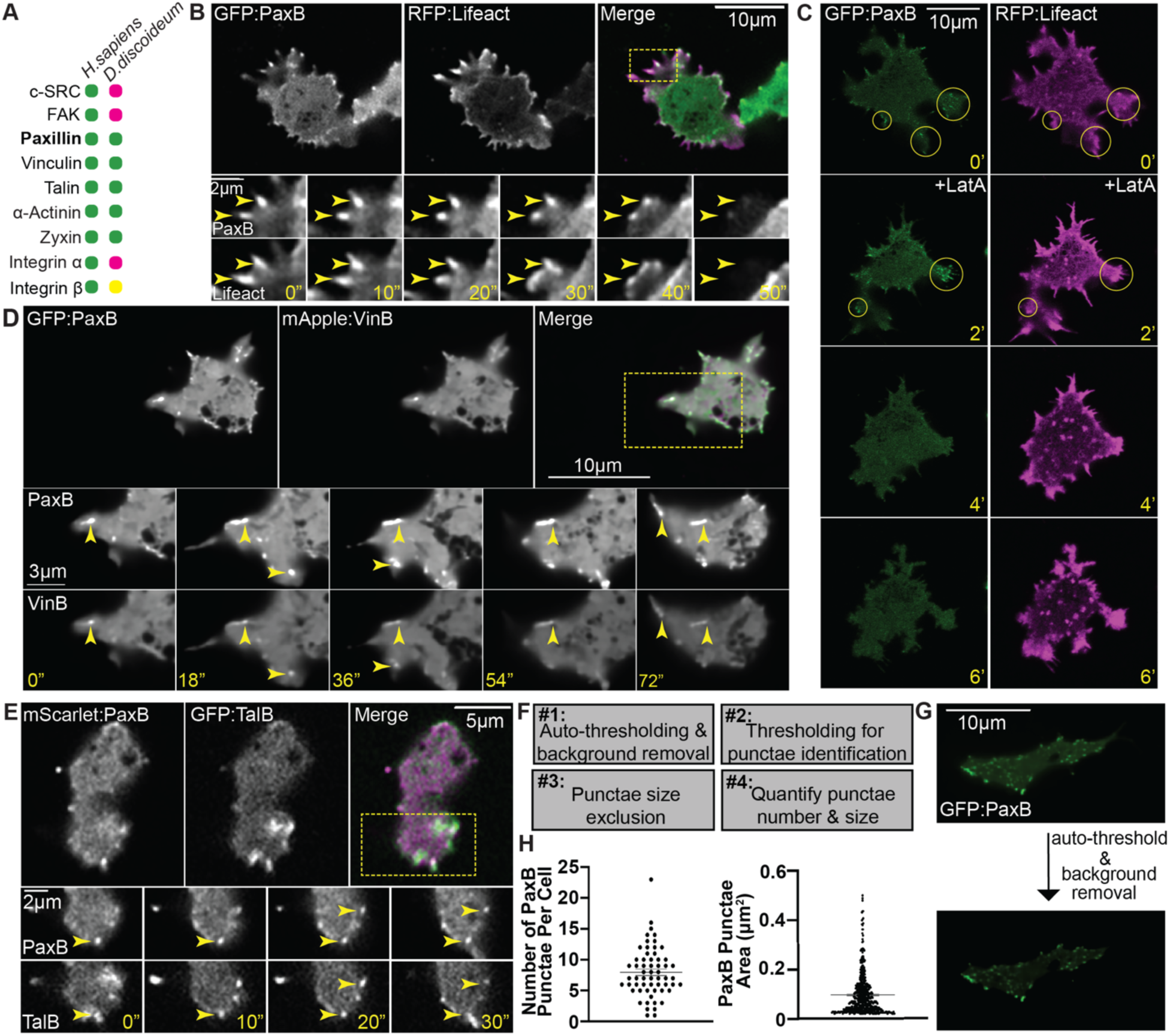
PaxillinB, the putative *Dictyostelium discoideum* homologue of the core adhesion scaffold Paxillin, forms small cell-substrate adhesion structures at the cell ventral surface. (A) Comparison of the presence and absence of core focal adhesion components between humans and the model Amoebozoa, *Dictyostelium discoideum*. Green indicates presence of putative homologue, yellow indicates potential putative homologue, and pink indicates no putative homologue. (B) Representative timelapse fluorescent confocal microscopy images of *paxB^−^*/act15/GFP:PaxB;RFP:Lifeact *Dictyostelium* cells. Yellow arrowheads shown on inset images show presence and disassembly of co-localizing PaxillinB and actin punctae over time. Time in seconds. (C) Representative timelapse fluorescent confocal microscopy images of PaxillinB (left) and actin (right) in act15/GFP:PaxB;RFP:Lifeact *Dictyostelium* cells before and after spike-in of 1 uM LatrunculinA (LatA) during imaging. Time in minutes. Spike-in of LatA indicated in image taken at 2-minute mark. Yellow circles in first two frames indicate sites of PaxB-positive punctae. (D) Representative timelapse fluorescent confocal microscopy images of *paxB^−^*/act15/GFP:PaxB;mApple:VinB. Yellow box in upper right panel indicates zoomed region shown in the lower panels. Lower panel images demonstrate co-localization of PaxillinB and VinculinB punctae, as indicated by yellow arrowhead. Time in seconds. **E)** Representative timelapse fluorescent confocal microscopy images of act15/GFP:TalB;mScarlet:PaxB. Yellow box in upper right panel indicates zoomed region shown in the lower panels. Lower panel images demonstrate co-localization of PaxillinB and TalinB punctae, as indicated by yellow arrowhead. Time in seconds. (F) Schematic showing workflow of image processing pipeline for PaxillinB punctae identification and quantification. See Methods for details. (G) Representative images prior to (top) and after (below) application of whole cell auto-thresholding and background removal to increase signal to noise ratio for PaxillinB punctae quantification. (H) Quantification of PaxillinB punctae number per cell (left) and average area of individual PaxillinB punctae (right) using pipeline described in (F); n=59 cells for left graph and n=505 punctae for right graph. Mean +/− SEM.

### The *Dictyostelium* Paxillin homologue localizes to cell-substrate adhesions at the ventral surface

The *Dictyostelium* homologue of the focal adhesion scaffolding molecule Paxillin is PaxillinB. In mammalian systems, Paxillin is a key scaffolding protein involved in regulating focal adhesion dynamics and cell migration via interactions with other focal adhesion components such as FAK, c-SRC, Vinculin, and Talin (Lopez-Colome et al., 2017; Schaller and Parsons, 1995; Turner et al., 1990). Initial work using PaxillinB overexpression in *Dictyostelium* suggests that PaxillinB localizes to ventral surface foci (Bukharova et al., 2005; Nagasaki et al., 2009; Patel et al., 2008; Pribic et al., 2011) reminiscent of mammalian focal adhesions (Beningo et al., 2001; Hu et al., 2014; Xue et al., 2023). Furthermore, knockout of PaxillinB decreased cell adhesion using plate shaking assays (Bukharova et al., 2005; Nagasaki et al., 2009), suggesting that PaxillinB functions in cell adhesion to the underlying substrate. How or whether PaxillinB regulates cell-substrate adhesion dynamics during *Dictyostelium* cell migration, however, is unclear. Additionally, *Dictyostelium* lacks putative homologues for FAK and cSRC – which phosphorylate Paxillin to regulate adhesion dynamics in mammalian systems and are classically thought of as cognate binding partners (Bellis et al., 1997; Burridge et al., 1992; Mitra et al., 2005; Sachdev et al., 2009; Schaller and Schaefer, 2001) – suggesting a unique and potentially novel mechanism for Paxillin regulation. Thus, we investigated the role of PaxillinB at *Dictyostelium* cell-substrate adhesions during migration.

We first sought to visualize PaxillinB localization with high-resolution microscopy approaches. Previous research examining PaxillinB overexpression in a *paxB^−^* background suggests overexpression does not perturb PaxillinB function (Bukharova et al., 2005). We overexpressed GFP-tagged PaxillinB under an actin15 promoter in a *paxB^−^* background. Imaging of fluorescently-tagged PaxillinB using spinning disc confocal microscopy in the *paxB^−^* background showed localization of PaxillinB to ventral surface foci (Figure 2B and Video S1), in line with previous imaging studies (Bukharova et al., 2005; Nagasaki et al., 2009; Patel et al., 2008; Pribic et al., 2011). Consistent with these structures being cell-substrate adhesions, we used total internal reflection fluorescence (TIRF) microscopy to determine that the PaxillinB-positive foci formed at the ventral surface of *Dictyostelium* cells (Figure S1A and Video S2). Furthermore, the PaxillinB-positive foci co-localized with actin, marked by RFP:Lifeact, at dynamic structures at the leading edge of the cell (Figure 2B and Video S1). Perturbation of actin polymerization via LatrunculinA spike-in, which is commonly used to perturb cell-substrate adhesions (Cai et al., 2000), led to ablation of PaxillinB foci shortly after spike-in (Figure 2C and Video S3), suggesting PaxillinB localization is actin-dependent. We also tested whether PaxillinB co-localizes with other *Dictyostelium* cell-substrate adhesion component homologues. Knockout of the two *Dictyostelium* Talin homologues, TalinA and TalinB, has been shown to perturb PaxillinB localization (Tsujioka et al., 2008). Furthermore, *Dictyostelium* possesses at least one homologue of Vinculin, VinculinB, which has been implicated in *Dictyostelium* cell motility and shown to bind to other known cytoskeletal and focal adhesion protein homologues (Huber and O’Day, 2012; Nagasaki et al., 2009). Therefore, we examined co-localization of PaxillinB with Talin and Vinculin molecules and found that PaxillinB co-localized with VinculinB and colocalized with TalinB at ventral surface foci during migration (Figure 2, D and E and Video S4 and S5). Altogether, these data suggest PaxillinB localizes to dynamic cell-substrate adhesions in *Dictyostelium*.

Focal adhesion size regulates adhesion-based migration in mesenchymal cells. Increased focal adhesion size correlates with increased migration speed until the adhesion size reaches a threshold at which point the speed decreases (Kim and Wirtz, 2013). Thus, we next sought to quantify the size as well as the number of PaxillinB-positive adhesions. We developed a semi-automated image processing pipeline using ImageJ to quantify single timepoint images of individual cells expressing GFP:PaxillinB. This pipeline allows us to increase the signal-to-noise ratio and identify and quantify individual PaxillinB-positive adhesions in an unbiased manner (Figure 2, F and G). Using this tool, we observed a wide range of Paxillin-positive adhesion number per cell ranging from one to twenty-three, with an average area of ∼0.096 microns^2^ (Figure 2H). The average size of these PaxillinB-positive adhesions is significantly smaller than previously described size criteria for focal adhesions in mesenchymal Metazoan systems, which are typically 0.6 microns^2^ (Gardel et al., 2010; Han et al., 2021; Han et al., 2015). Indeed, PaxillinB adhesion size is more reminiscent of focal adhesions visualized in physiological contexts (∼0.1-0.2 microns^2^ for “curved” adhesions) (Zhang et al., 2023) and for mesenchymal zebrafish melanoma cells *in vivo* (∼0.3 microns^2^) (Xue et al., 2023)), as well as nascent focal contacts in mammalian cells (0.24 microns^2^) (Han et al., 2021; Han et al., 2015). In mammalian models, substrate composition has been shown to influence focal adhesion formation and dynamics (Grudtsyna et al., 2023; Kato and Mrksich, 2004; Pelham and Wang, 1997; Schneider and Burridge, 1994; Seo et al., 2011; Zhou et al., 2017), so we next investigated the effect of different substrates on the number and size of PaxillinB-positive adhesions. Though homology analyses suggest *Dictyostelium* cells lack clear homologues of ECM proteins traditionally associated with focal adhesion formation (Hynes, 2012; Linden, 2022), previous work has shown that altering the underlying substrate influences *Dictyostelium* cell adhesion and migration (Decave et al., 2002; Loomis et al., 2012; McCann et al., 2014). Therefore, we plated cells on glass, bovine serum albumin (BSA), or poly-L-lysine (PLL). We found that cells plated on BSA formed significantly fewer and smaller PaxillinB punctae compared to cells on glass, while cells on PLL formed bigger, but not more, PaxillinB punctae than cells on glass, suggesting that the substrate impacts PaxillinB punctae formation and size (Figure S1, B-D). Together, these data suggest *Dictyostelium* is a non-Metazoan organism capable of forming PaxillinB-positive adhesion structures.

### Knockout of PaxillinB reduces cell adhesive capacity

Next, we sought to determine the effect of PaxillinB knockout on *Dictyostelium* cell adhesive capacity. First, we used differential interference contrast microscopy (DIC) to quantify cell spreading – often used as an indicator of cell adhesive capability – of wild-type and *paxB^−^* cells (Figure 3A). We found that knockout of PaxillinB led to significantly decreased cell spreading (Figure 3B), suggesting reduced cell adhesive capacity. To assay cell adhesion strength more directly, we used an orthogonal microfluidics system (Figure 3C) to test how increasing shear flow forces affected the adhesion of wild-type and *paxB^−^* cells to the substrate. We determined that *paxB^−^* cells trended toward a higher detachment rate when exposed to increasing fluid flow rates compared to wild-type cells (Figure 3D and Figure S1E). These data suggest PaxillinB promotes *Dictyostelium* cell adhesion to the underlying substrate.

**Figure 3:**
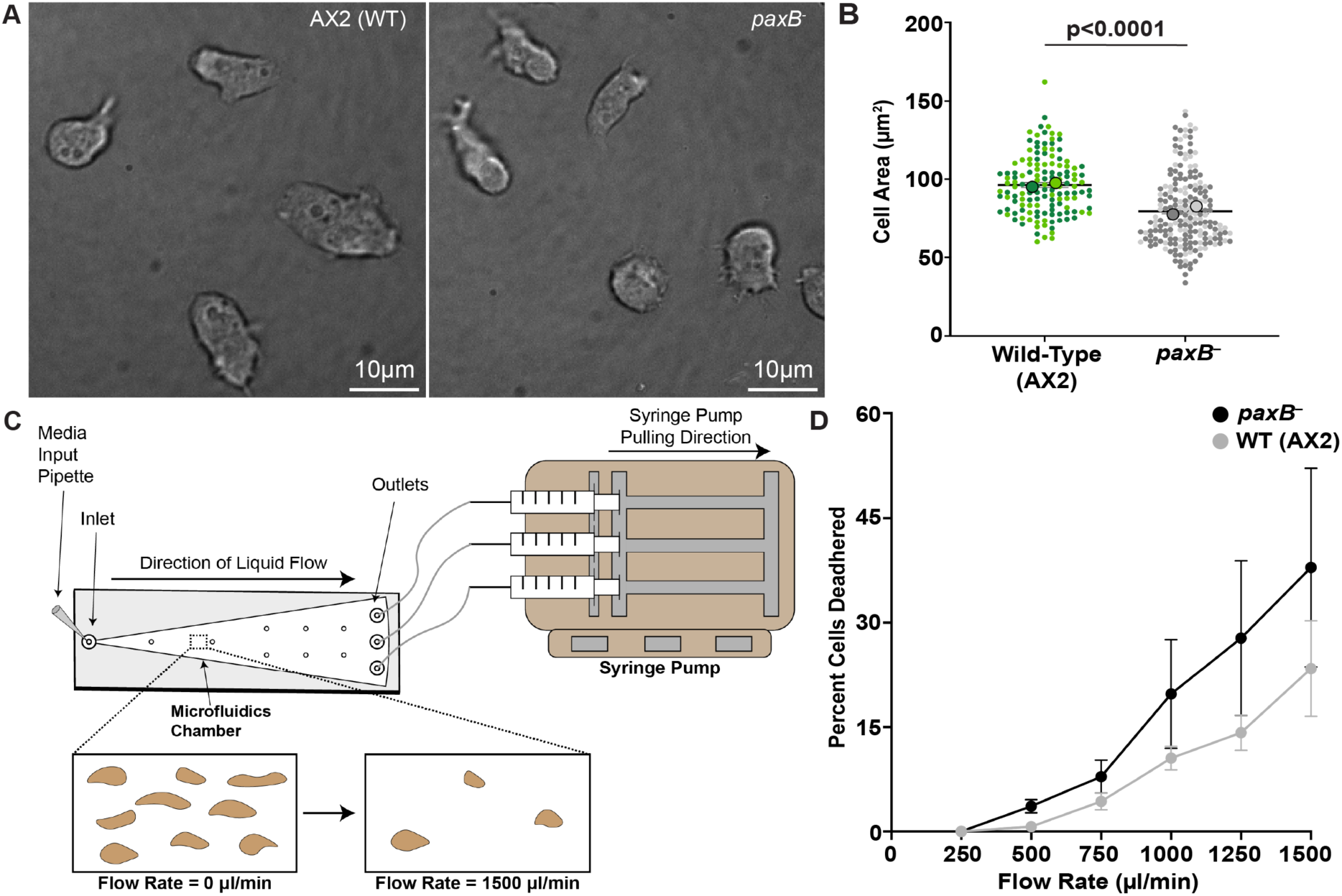
PaxillinB regulates *Dictyostelium* cell-substrate adhesion capability. (A) Representative differential interference contrast (DIC) images of wildtype and *paxB^−^ Dictyostelium* cells that were used to quantify cell spreading. (B) Average whole cell area of wildtype and *paxB^−^ Dictyostelium* cells. Student t-test. n=136 (wild-type) and 202 (*paxB^−^*) cells across n=2 biological replicates. (C) Cartoon schematic of the microfluidics assay used to test adhesive capability (see Methods). For each individual assay, cells were imaged in the same region of the device (small square in schematic) to ensure exposure to similar shear flow stress across replicates. (D) Quantification of wildtype (AX2) and *paxB^−^*cell detachment at increasing flow rates. N=4 experiments; mean +/− SEM; Student t-test. Only comparisons that are statistically significant are shown.

### C-terminal LIM domains are required for PaxillinB localization

We next determined which domains are required for PaxillinB localization and function at *Dictyostelium* cell-substrate adhesions. Previous work (Bukharova et al., 2005) used pairwise sequence alignments of *Dictyostelium* and human Paxillin molecules to show conservation of N-terminal leucine-rich LD motifs and C-terminal LIM (**L**in-11, **I**sl-1 and **M**ec-3) domains, but not of the Tyrosine 31 (Y31) and Tyrosine 118 (Y118) residues that are phosphorylated by upstream kinases, FAK and c-SRC to regulate focal adhesion turnover and cell migration (Bellis et al., 1997; Burridge et al., 1992; Mitra et al., 2005; Sachdev et al., 2009; Schaller and Schaefer, 2001). We sought to build on this work by using multiple sequence alignments of Paxillin molecules from various organisms to identify Paxillin features that are conserved across evolutionary space and likely to be fundamental aspects of cell-substrate adhesion function and regulation across the species tree of life. Furthermore, we used domain architecture and predicted structural homology tools (McGinnis and Madden, 2004; Potter et al., 2018; van Kempen et al., 2024) utilized for our bidirectional homology pipeline (Figure 1) to provide higher sensitivity during comparative analyses.

Comparisons of the structure, domain, and sequence architecture of Paxillin molecules across several organisms show conservation across the N-terminal LD motifs and C-terminal LIM domains (Figure 4A). Quantitative homology analyses between human and *Dictyostelium* Paxillin molecules showed a higher degree of structural and sequence similarity between the C-terminal LIM domains of the two proteins versus either the N-termini alone or the full-length protein (Figure 4B). While vertebrate Paxillin proteins possess five LD motifs at the N-terminus, which are binding sites of other focal adhesion proteins (Lopez-Colome et al., 2017), non-vertebrate Paxillin molecules possess varying numbers of LD motifs, such as the four LD motifs in *Dictyostelium* PaxillinB (Figure 4A). The sequences surrounding the LD motifs, however, are not well-conserved. Thus, we examined these specific motifs closer, focusing on residues important for the FAK-Paxillin interaction axis in mammalian systems. Using multiple sequence alignments, we observed Paxillin Y31 and Y118 residues are limited to Vertebrate and Metazoan Paxillin molecules, respectively, as none of the non-Metazoan Paxillin proteins, including *Dictyostelium* PaxillinB, possessed these residues (Figure 4C). We also used multiple sequence alignments to evaluate conservation of key aspartic acid and arginine residues in the LD2 motif necessary for FAK binding to Paxillin molecules (Brown et al., 1996). While the aspartic acid residue was well conserved, the arginine residue was only found in vertebrate Paxillin proteins and was not substituted with another basic amino acid (Figure 4D). These data, paired with the lack of a FAK homologue (Figures 1 and 2A) or any other compensatory tyrosine kinase in *Dictyostelium* (Goldberg et al., 2006), suggest that the FAK-Paxillin binding axis and phosphorylation activity are not conserved in *Dictyostelium*. Furthermore, these observations suggest that Paxillin activity is not regulated by tyrosine phosphorylation in *Dictyostelium,* as it is in Metazoan systems.

**Figure 4:**
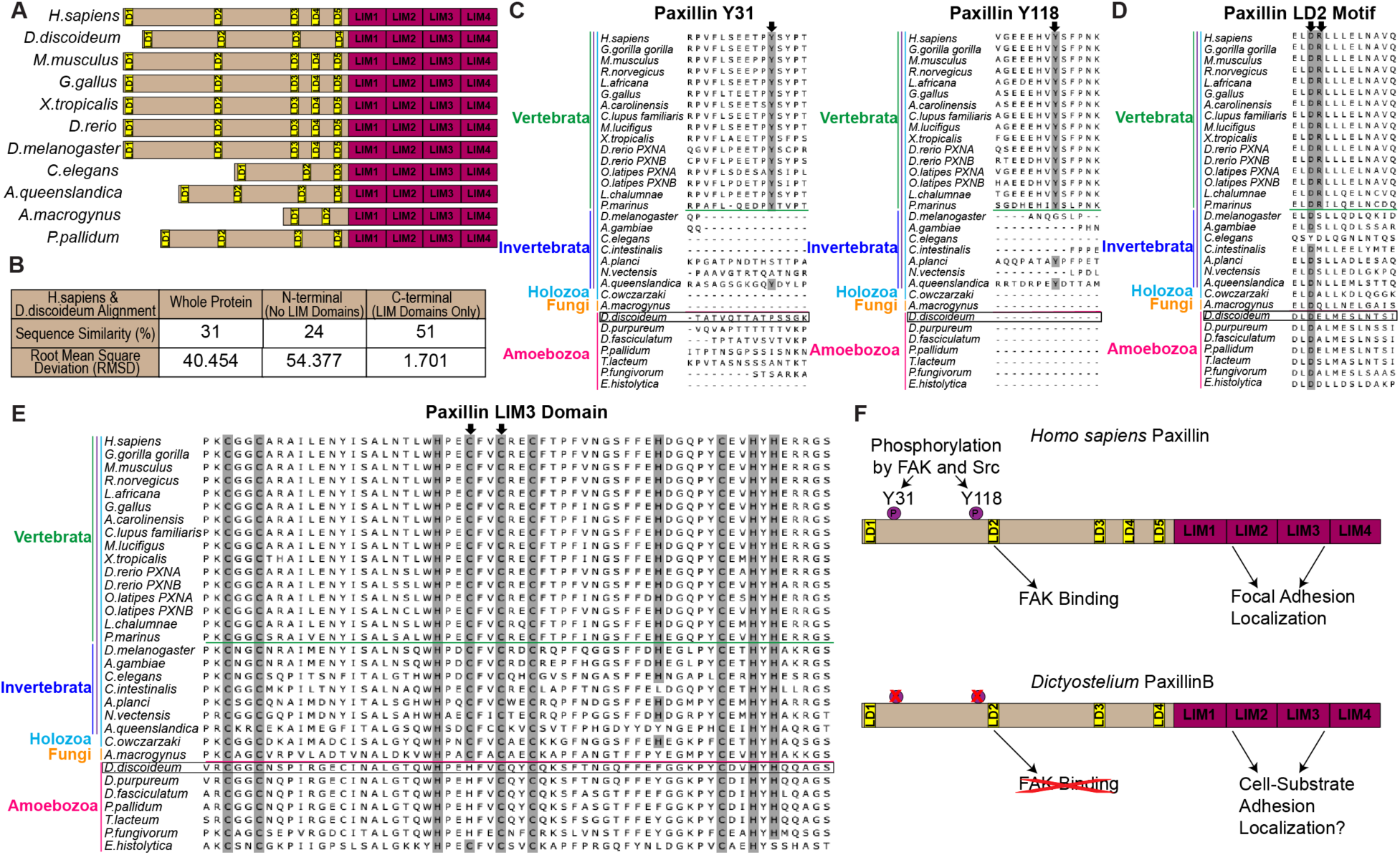
PaxillinB has conserved C-terminal LIM domains but does not exhibit conservation of key N-terminal residues. (A) Schematic of Paxillin protein structure/domains across representative organisms. (B) Quantification of sequence (percent similarity) and structural (using root mean square deviation (RMSD) of Alphafold predicted structures) conservation between aligned human and *Dictyostelium* Paxillin molecules. (C) Multiple Sequence Alignments (MSAs) of Paxillin Tyrosine 31 and 118, phosphorylation sites associated with focal adhesion turnover in Metazoans. Black arrowheads point to the tyrosine residues of interest. (D) MSAs of Paxillin LD2 motifs, a known binding site for FAK in Metazoan adhesions. Black arrowheads indicate aspartic acid and arginine residues shown to be critical for FAK binding in Metazoan literature. (E) MSAs of Paxillin LIM3 domains, which has been shown to be necessary for Paxillin localization to Metazoan focal adhesions. Cysteine and histidine residues are highlighted in gray, respectively. Black arrowheads indicate conservation of either a cysteine or histidine at key sites corresponding to the zinc-coordinating residues required for Paxillin localization to focal adhesions, as previously described (Brown et al., 1996). (F) Schematic showing comparison of presence and absence of conserved adhesion-specific Paxillin domains between human and *Dictyostelium* Paxillin molecules.

We next focused on the conserved C-terminal LIM domains involved in focal adhesion functions (Figure 4, A and B). We performed phylogenetic analyses of the Paxillin LIM domains across Eukaryotic species, along with the LIM domains of other LIM domain-containing proteins. These analyses showed that the LIM domains of Amoebozoan Paxillin molecules, including *Dictyostelium* PaxillinB, cluster with other Paxillin molecules as well as Leupaxin, a vertebrate-specific member of the Paxillin superfamily (Figure S2A). These data suggest that *Dictyostelium* PaxillinB LIM domains are evolutionarily similar to the LIM domains of other Paxillin molecules. Previous functional analyses of Paxillin LIM domains suggest the LIM3 domain, specifically two zinc-coordinating cysteine residues found in a zinc-finger motif in the domain, is necessary for localization of Paxillin to focal adhesions in mammalian cells (Brown et al., 1996). Multiple sequence alignments of only the LIM3 domain of Paxillin proteins suggest that these cysteine residues are well conserved across species, with Amoebozoan Paxillin molecules – including *Dictyostelium* PaxillinB – possessing an acceptable histidine substitution (Cassandri et al., 2017) at one of these residues (Figure 4E). A closer examination of this histidine residue in the Alphafold-predicted structure of *Dictyostelium* PaxillinB suggests the zinc-coordinating pocket is conserved relative to the predicted human Paxillin structure (Figure S2B). Altogether, these data show *Dictyostelium* PaxillinB conservation of the C-terminal LIM domains involved in localization to focal adhesions, but not the N-terminal residues involved in FAK binding and phosphorylation (Figure 4F).

### Perturbation of PaxillinB LIM domains decreases adhesion localization and increases cell migration speed and displacement

The Paxillin LIM3 domain has previously been shown to be required for Paxillin localization to focal adhesions in mammalian cells (Brown et al., 1996; Brown et al., 1998; Brown and Turner, 2002; Ripamonti et al., 2021). Due to the conservation of the LIM3 domain in *Dictyostelium* PaxillinB at the sequence and predicted protein folding level, we next tested for functional homology of this domain. We generated cells in which truncated GFP-tagged PaxillinB molecules were expressed in the *paxB^−^* background (Figure 5A) and evaluated adhesion localization. Cells expressing either wild-type PaxillinB (GFP:PaxB-WT) or a truncated PaxillinB only lacking the LIM4 domain (GFP:PaxB-ΔL4) but still containing all of the other LIM domains, showed PaxillinB localization to adhesions (Figure 5B). To our surprise, however, cells expressing truncated PaxillinB proteins that lacked the LIM3 domain, but had at least one other LIM domain (GFP:PaxB-ΔL3, GFP:PaxB-ΔL34 and GFP:PaxB-ΔL234), still exhibited PaxillinB punctae formation (Figure 5B), suggesting that PaxillinB was still able to localize to cell-substrate adhesions in the absence of LIM3. Quantification of PaxillinB-positive adhesions revealed that the absence of the LIM3 domain, either by itself or with the absence of other LIM domains, led to a significant decrease in the average number of PaxillinB-positive adhesions formed per cell (Figure 5C), with no significant impact on punctae size (Figure S3A). Deletion of all LIM domains (GFP:PaxB-ΔLIMs) led to full disruption of PaxillinB-positive adhesion formation (Figure 5, B and C). These data suggest that the presence of at least one LIM domain is required for PaxillinB localization to adhesions, with the LIM3 domain still important, but not solely necessary, for efficient localization of PaxillinB to adhesions. We next tested whether removing the LIM domains affected cell adhesive capability to the substrate. We used our previously described microfluidics system to compare the adhesion of GFP:PaxB-WT cells vs GFP:PaxB-ΔLIMs cells to the substrate. As expected, GFP:PaxB-ΔLIMs cells trended toward a higher detachment rate compared to GFP:PaxB-WT cells (Figures 5D and S3B), suggesting PaxillinB localization to adhesions via the LIM domains is important for adhesive capability.

**Figure 5:**
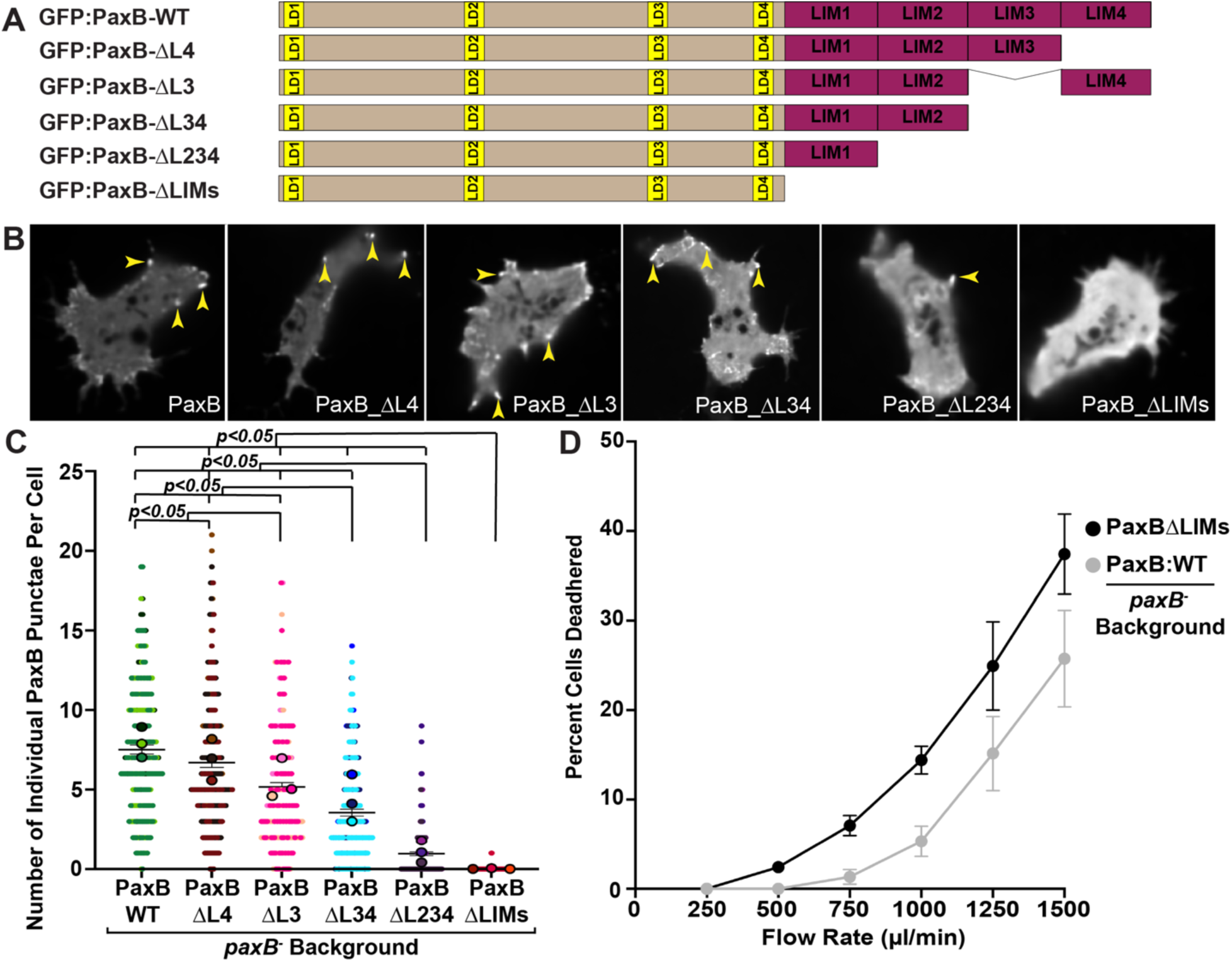
Perturbation of PaxillinB LIM domains decreases PaxillinB localization to cell-substrate adhesions. (A) Schematic demonstrating wild-type and LIM domain truncated PaxillinB molecules that were overexpressed in *paxB^−^ Dictyostelium* cells. (B) Representative images of timelapse fluorescent confocal microscopy of *paxB^−^ Dictyostelium* cells overexpressing the PaxillinB molecules shown in Figure 5A. Yellow arrowheads point to PaxillinB punctae. (C) Quantification of number of PaxillinB punctae per cell in *paxB^−^ Dictyostelium* cells overexpressing PaxillinB molecules shown in Figure 5A. n = 211 (PaxB-WT), 209 (PaxB-ΔL4), 203 (PaxB-ΔL3), 211 (PaxB-ΔL34), 210 (PaxB-ΔL234), and 213 (PaxB-ΔLIMs) cells across n=3 biological replicates per cell line. Mean +/− SEM; Kruskal-Wallis Test. (D) Quantification of cell detachment of *paxB^−^ Dictyostelium* cells overexpressing either GFP:PaxB-WT or GFP:PaxB-ΔLIMs proteins to compare cell adhesion capability. N=3 experiments; mean +/− SEM; Student t-test. For all graphs, only comparisons that are statistically significant are shown.

Due to the decreased number of PaxillinB-positive adhesions in PaxillinB LIM domain mutants, we next sought to test the effects of these truncations on migration capability. Previous work has shown that truncation of the Paxillin C-terminal LIM domains significantly decreases both cell migration speed and persistence in mammalian cells (Sero et al., 2011). We quantified *Dictyostelium* cell migration speed and mean square displacement, which is a measure of cell displacement and relative distance explored (Gorelik and Gautreau, 2014; Loosley et al., 2015), during random migration. When expressing wildtype Paxillin in the *paxB^−^* background, we surprisingly found no difference in cell migration speeds compared to *paxB^−^* cells alone (comparing “PaxB WT” to “GFP empty” in Figure 6A). Unexpectedly, cells expressing truncated PaxillinB proteins demonstrated *increased* average cell speed and mean square displacement relative to those expressing wildtype PaxillinB (Figure 6, A and B), and this change in migration speed was observed regardless of whether we truncated one or multiple C-terminal LIM domains. Furthermore, correlation analyses of cell migration speeds and PaxillinB expression levels of individual cells confirmed changes in cell migration speed was not an artifact of differential PaxillinB expression levels (Figure S4). These results suggest that first, each PaxillinB LIM domain regulates *Dictyostelium* random cell migration; second, these results show that, surprisingly, decreasing the number of PaxillinB-positive adhesions increases cell migration speed and displacement. Due to this unexpected observation, we next asked whether this phenotype was recapitulated when *Dictyostelium* cells underwent directed cell migration toward a chemotactic cue in a process called chemotaxis. *Dictyostelium* undergo cAMP chemotaxis in response to starvation as cells transition to multicellularity, and this transition is accompanied by increased migration velocity (Bozzaro, 2019). We used a chemotaxis assay (Figure 6C) which allows for robust visualization and analysis of *Dictyostelium* cells undergoing directed cell migration towards a chemoattractive cAMP source. To ensure that the cAMP gradient was properly established, we only used datasets in which >80% of wildtype cells migrated directionally toward cAMP using a spider plot analysis (Figure 6D). Consistent with our random cell migration results, we found that during chemotaxis, cells expressing GFP:PaxB-ΔL34 trended toward faster migration speeds than cells expressing PaxB-WT, and cells expressing GFP:PaxB-ΔLIMs exhibited significantly increased cell migration speeds compared to wildtype (Figure 6E). Together, these results suggest that removing PaxillinB LIM domains decreases PaxillinB localization to cell-substrate adhesions, which leads to increased migration speeds during both random and directed migration.

**Figure 6:**
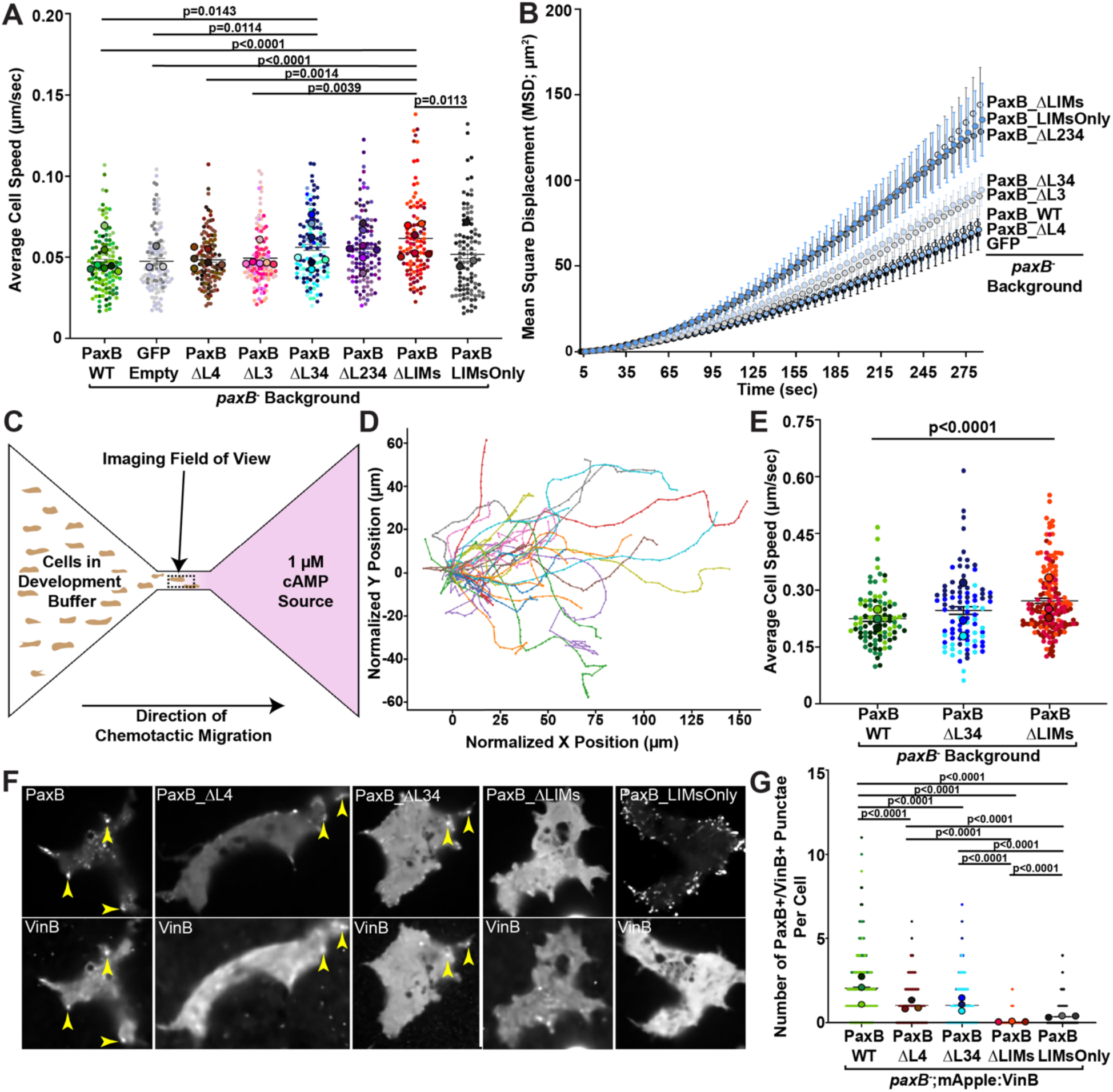
Cells with decreased number of PaxillinB-VinculinB double positive cell-substrate adhesions exhibit increased migration speeds. (A and B) Quantification of average cell speed (A) and mean square displacement (B) of randomly migrating *paxB^−^ Dictyostelium* cells overexpressing PaxillinB molecules shown in Figure 5A or a GFP Empty control. For both A) and B), n = 113 (PaxB-WT), 122 (GFP), 119 (PaxB-ΔL4), 108 (PaxB-ΔL3), 119 (PaxB-ΔL34), 111(PaxB-ΔL234), 108 (PaxB-ΔLIMs), and 103 (PaxB-LIMsOnly) cells across n=3-6 biological replicates per cell line. Kruskal-Wallis Test. Mean +/− SEM. (C) Schematic showing setup of cAMP chemotaxis assay. Cells in development buffer are plated to the left of the slide and the cAMP source is on the right. The dotted black box indicates the imaging field of view. (D) A representative spider plot used to determine whether cAMP gradients were appropriately generated, with each cell’s migration trajectory as a separate color. (E) Quantification of average cell speed during chemotactic migration; n = 93 (PaxB-WT), 90 (PaxB-ΔL34) and 161 (PaxB-ΔLIMs) across n=3 biological replicates per cell line. Brown-Forsythe and Welch ANOVA tests. Mean +/− SEM. (F) Representative images of timelapse fluorescent confocal microscopy of *paxB^−^ Dictyostelium* cells overexpressing the PaxillinB molecules indicated in each image. Yellow arrowheads point to PaxillinB-positive and VinculinB-positive punctae (PaxB+;VinB+). (G) Quantification of the total number of PaxB+;VinB+ punctae per cell in each cell line; n = 745 (PaxB-WT), 646 (PaxB-ΔL4), 676 (PaxB-ΔL34), 593 (PaxB-ΔLIMs), and 552 (PaxB-LIMsOnly) cells across n=3 biological replicates per cell line. For all graphs, only comparisons that are statistically significant are shown.

### Migration phenotypes in PaxillinB LIM domain mutants are not due to a transition to amoeboid migration or changes in PaxillinB dynamics

Due to the surprising observation that decreasing the number of Paxillin-positive adhesions led to increased cell migration speeds, we first asked whether this increase in cell migration speed was due to a transition from an adhesion-based form of migration to an adhesion-independent amoeboid migration, which is often associated with faster migration speed (Alexandrova et al., 2020; Pankova et al., 2010). A hallmark of amoeboid migration is increased cortical contractility and accumulation of actin at the rear of the cell (Alexandrova et al., 2020; Callan-Jones and Voituriez, 2016). Thus, we quantified the ratio of actin density at the rear versus the front of migrating cells. Quantification of this ratio showed no significant enrichment of actin at the rear versus the front of *Dictyostelium* cells expressing truncated PaxillinB compared to those expressing wildtype PaxillinB (Figure S5A), suggesting that *Dictyostelium* cells are not transitioning to adhesion-independent amoeboid motility in the Paxillin LIM domain mutants.

We next asked if changes in the dynamics of PaxillinB-positive adhesions could explain the cell migration phenotype. We quantified the lifetime of PaxillinB-positive adhesions in *Dictyostelium* cells expressing either GFP:PaxB-WT or the truncated GFP:PaxB-ΔL34 protein. The GFP:PaxB-ΔL34 line was chosen as these cells exhibited significant perturbation of PaxillinB localization and increased cell migration speed, but still formed sufficient numbers of PaxillinB-positive adhesions for quantification. We did not observe differences in PaxillinB-positive adhesion lifetime in cells expressing GFP:PaxB-ΔL34 compared to cells expressing GFP:PaxB-WT (Figure S5B). Altogether, these data suggest that the increased cell migration speed and displacement phenotypes in the PaxillinB LIM domain mutants are not a result of either a transition to adhesion-independent amoeboid motility or a change in PaxillinB-positive adhesion dynamics.

### The number of PaxillinB and VinculinB positive cell-substrate adhesions correlates with cell migration speed and displacement

We next decided to further investigate if the increased cell migration speed and displacement observed in the Paxillin LIM domain mutants were due to perturbation of cell-substrate adhesion number. To rigorously identify and quantify cell substrate adhesions, we used VinculinB as a secondary marker and defined “bona fide” cell-substrate adhesions as adhesions that were positive for *both* VinculinB and PaxillinB. VinculinB colocalizes with PaxillinB at *Dictyostelium* cell-substrate adhesions (Figure 2D) and previous work in mammalian cells suggests Paxillin and Vinculin are binding partners at focal adhesions (Brown et al., 1996; Turner et al., 1990). We generated *paxB^−^* cells that co-expressed mApple-tagged VinculinB and GFP-tagged wild-type or truncated PaxillinB. Quantification of PaxillinB+/VinculinB+ cell-substrate adhesions revealed a decreased number of cell substrate adhesions in cells expressing GFP:PaxB-ΔL4 and GFP:PaxB-ΔL34 relative to cells expressing GFP:PaxB-WT (Figure 6, F and G). These results suggest that perturbing PaxillinB LIM domains decreases cell-substrate adhesion number. Combined with our cell migration results, these data suggest that decreasing cell-substrate adhesion number promotes *Dictyostelium* cell migration speed and displacement.

To test our model, we next asked whether *increasing* cell-substrate adhesion number would then *decrease* cell migration speed and displacement. We generated a PaxillinB construct containing only the LIM domains (GFP:PaxB-LIMsOnly), which we predicted would promote PaxillinB localization to cell-substrate adhesions. Expression of GFP:PaxB-LIMsOnly led to a dramatic increase in the number, but not size, of PaxillinB-positive adhesions (Figures 6F, S6A and B). To our surprise, however, the dramatic increase in PaxillinB-positive punctae number was concomitant with diffuse VinculinB localization, suggesting failed VinculinB localization to PaxillinB-positive adhesions upon truncation of the N-terminal half of PaxillinB (Figure 6F). Indeed, when we quantified the number of PaxillinB+/VinculinB+ cell-substrate adhesions in GFP:PaxB-LIMsOnly expressing cells, we found that the number of PaxillinB+/VinculinB+ cell-substrate adhesions was low and even lower than the observed number in PaxillinB LIM domain mutants (Figures 6G, S6C-E). Given this surprising finding, we then hypothesized that the GFP:PaxB-LIMsOnly cells would have increased cell migration speeds due to decreased number of PaxillinB+/VinculinB+ cell-substrate adhesions. Consistent with this hypothesis, GFP:PaxB-LIMsOnly cells exhibited increased cell migration speed and displacement compared to wildtype PaxillinB, at a rate similar to the Paxillin LIM domain mutants (Figure 6, A and B). Altogether, these results suggest a model by which the formation of cell substrate adhesions, as marked by PaxillinB+/VinculinB+, specifically dictate cell migration speed in an inverse relationship, with *decreased* number of cell-substrate adhesions leading to *increased* cell migration speeds (Figure 7).

**Figure 7:**
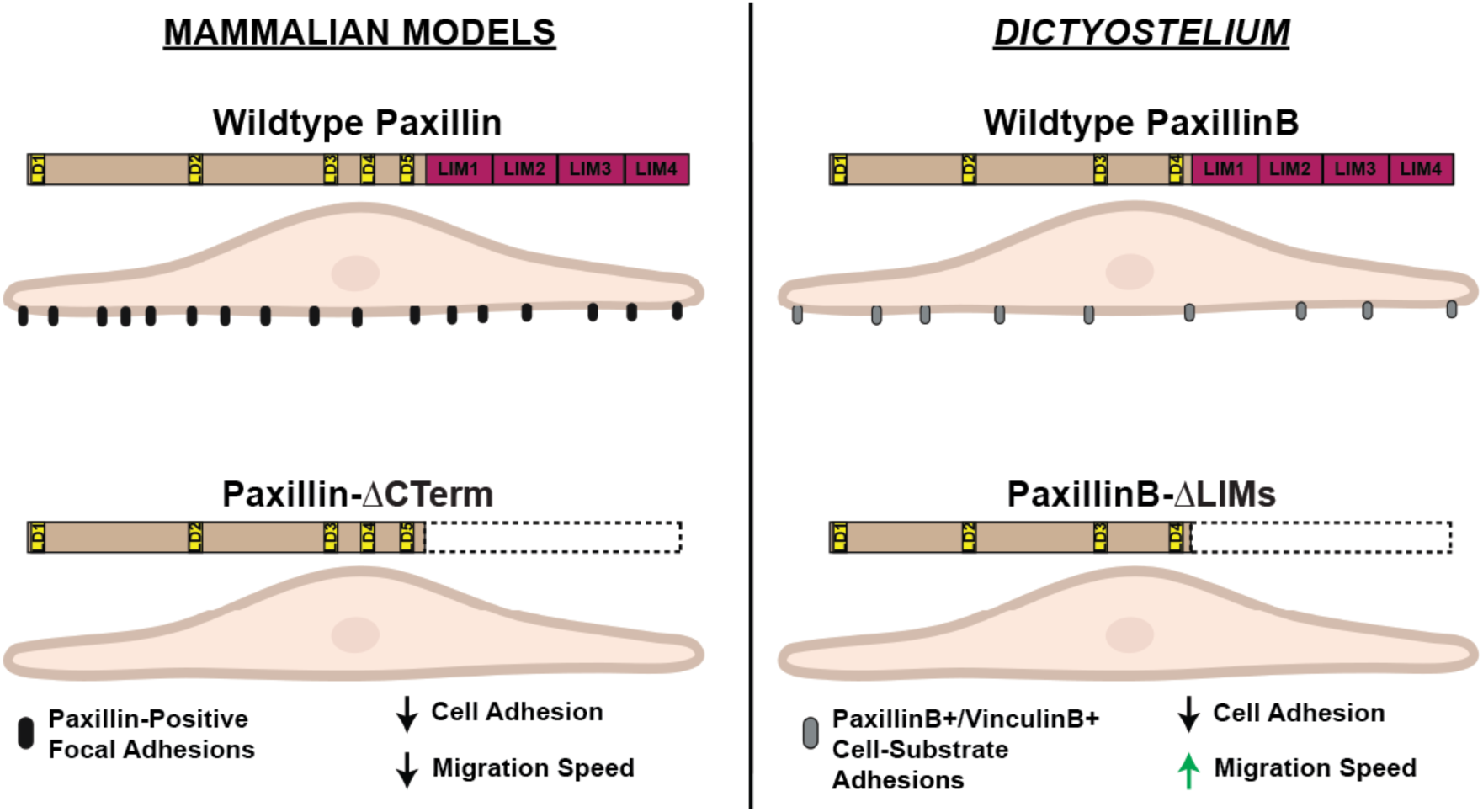
Proposed Model – PaxillinB localizes to *Dictyostelium* cell-substrate adhesions that serve as molecular “brakes” to promote cell adhesion and regulate cell migration speed. We speculate that cell-substrate adhesions are used by *Dictyostelium* to enable efficient adhesion to the underlying substrate and reduce cell migration speed, acting as a molecular “brake”. These cell-substrate adhesions are double-positive for PaxillinB and VinculinB, with perturbation of Paxillin localization via LIM domain truncation decreasing cell adhesion and *increasing* cell migration. This latter phenotype is in direct contrast to mammalian models to the left, where truncation of the Paxillin LIM domains specifically has been shown to decrease Paxillin localization and cell adhesion (Brown et al., 1996; Brown et al., 1998; Brown and Turner, 2002; Ripamonti et al., 2021) but cell migration speed is *decreased* (Sero et al., 2011).

## DISCUSSION

Cell-substrate adhesion composition, function, and regulation during cell migration is largely under-studied in non-Metazoan organisms. Work suggesting various non-Metazoan species can form cell-substrate adhesions despite possessing only some conserved core focal adhesion molecules (Fierro Morales et al., 2022; Sebé-Pedrós et al., 2010), however, makes non-Metazoans intriguing models for studying how cell-substrate adhesions have evolved in composition and function to regulate processes such as cell adhesion and migration. In this work, we investigate cell-substrate adhesion formation and regulation during cell migration in the non-Metazoan *Dictyostelium discoideum.* Using this model, we have uncovered an unexpected relationship between cell-substrate adhesion formation and cell migration. We first found that *Dictyostelium* cells form small, dynamic cell-substrate adhesions positive for PaxillinB and VinculinB, the *Dictyostelium* homologues of core focal adhesion proteins Paxillin and Vinculin, respectively. Consistent with regulation in mammalian cells, the conserved C-terminal LIM domains are required for PaxillinB localization to *Dictyostelium* cell-substrate adhesions. We also found that cells with reduced PaxillinB adhesion localization exhibited reduced adhesion to substrates under shear forces. However, when we quantified cell migration, we unexpectedly found that reducing cell-substrate adhesion number led to *increased* migration speeds and displacement. These findings are in direct contrast to findings in mammalian cells and lead us to speculate on a fundamentally different function for focal adhesion-like structures during cell migration in organisms evolutionarily distant from Metazoans (Figure 7).

Previous work in mammalian cells has shown that perturbing the Paxillin LIM3 domain reduces Paxillin localization focal adhesions and cell adhesion to the ECM (Brown et al., 1996; Brown et al., 1998; Brown and Turner, 2002; Ripamonti et al., 2021). Consistent with focal adhesion formation being required for cell migration, truncation of the C-terminal LIM domains of Paxillin alone *decreases* mammalian cell migration speed (Sero et al., 2011). Our data in *Dictyostelium* show the opposite, as decreasing the number of PaxillinB+/VinculinB+ cell-substrate adhesions through PaxillinB LIM domain truncation *increases Dictyostelium* cell migration speed. This key difference in the relationship between cell-substrate adhesion formation and migration speed when comparing mammalian and *Dictyostelium* cell-substrate adhesions provides an interesting window into the functional evolution of cell-substrate adhesions. We put forward a model whereby the ancestral function for cell-substrate adhesions may have been for adhesion to a substrate, and that these cell-substrate adhesions were then co-opted for enabling efficient migration. Below we summarize data that supports this model.

In mammalian systems, focal adhesions are required for a cell to adhere and migrate via adhesion-based migration (Barnhart et al., 2015; Case et al., 2015; Case and Waterman, 2015; Doyle et al., 2015; Gardel et al., 2010; Gupton and Waterman-Storer, 2006; Hu et al., 2014; Kanchanawong et al., 2010; Kim and Wirtz, 2013; Kowalewski et al., 2015; Kumari et al., 2024; Sero et al., 2011; Vicente-Manzanares and Horwitz, 2011; Yamaguchi and Knaut, 2022). Focal adhesion formation can lead to a range of migration phenotypes ranging from stationary cells (via formation of several large, long-lasting focal adhesions) (Angers-Loustau et al., 1999; Bear et al., 2000; Huttenlocher et al., 1996; Ilic et al., 1995; Kim and Wirtz, 2013; Ren et al., 2000; Smilenov et al., 1999) to highly migratory cells (via formation of dynamic and spatially regulated focal adhesions that undergo efficient turnover) (Beningo et al., 2001; Caillier et al., 2024; Gupton and Waterman-Storer, 2006; Kim and Wirtz, 2013; Lawson et al., 2012; Nayal et al., 2006; Palecek et al., 1997; Plotnikov et al., 2012; Xia et al., 2008), depending on factors such as adhesion size and number (Kim and Wirtz, 2013). Mammalian cells exhibit reduced cell migration speeds upon decreasing focal adhesion number because of an inability to adhere and pull on the ECM to propel the cell forward (De Pascalis and Etienne-Manneville, 2017; Gardel et al., 2010; Ntantie et al., 2018; Zhao et al., 2016). The pulling forces on the ECM are transduced into the cell via focal adhesions and the tension is maintained through the formation of stress fibers, which are bundled actin cables that associate with focal adhesions to promote adhesion maturation, mechanotransduction, and actin cytoskeletal reorganization (Choi et al., 2008; Kanchanawong et al., 2010; Oakes et al., 2012; Sun et al., 2020; Tojkander et al., 2012; Winkelman et al., 2020). Interestingly, *Dictyostelium* lack both actin stress fibers (Iwadate and Yumura, 2008b; Rubino et al., 1984) and clear ECM protein orthologues, with most ECM components arising after the divergence of Amoebozoa (Hynes, 2012; Linden, 2022). It is also unclear if *Dictyostelium* cells secrete ECM proteins for substrate adhesion while in a unicellular form. Indeed, it is unknown if the putative *Dictyostelium* beta integrin receptors, the Sib proteins (Cornillon et al., 2008; Cornillon et al., 2006), have cognate ECM substrates they attach to or if these surface glycoproteins are primarily involved in non-specific interactions such as Van der Waals forces, as previously proposed (Loomis et al., 2012). Given this, it is reasonable to speculate that *Dictyostelium* cells do not use cell-substrate adhesions for efficient cell migration via forces generated on an underlying substrate. Rather, *Dictyostelium* cells may form PaxillinB+/VinculinB+ cell-substrate adhesions as molecular “brakes” to regulate cell migration speed by increasing cell adhesion to the underlying substrate during migration. This hypothesis aligns with early observations in which *Dictyostelium* cells with increased number of actin-rich foci on the ventral surface exhibited decreased migration speed (Uchida and Yumura, 2004). An ability to migrate quickly and be able to form adhesion structures for “braking” purposes is relevant to *Dictyostelium-*specific processes such as the transition to multicellularity, where being able to briefly slow down and re-orient without “stalling” in response to changing environmental cues is important for finding other cells. With this proposed model, the lack of change in cell migration speed between *paxB^−^*cells and *paxB^−^* cells expressing a wildtype GFP:PaxillinB rescue (Figure 6A) may seem surprising, as we would expect the former to migrate faster. We posit, however, that these results may be due to fact that fully knocking out PaxillinB likely disrupts other processes besides formation of PaxillinB+/VinculinB+ cell-substrate adhesions that could, in turn, impair increased cell migration, though future work is needed to test this hypothesis.

The hypothesis that PaxillinB+/VinculinB+ cell-substrate adhesions serve as molecular “brakes” to regulate cell migration speed is further supported by the increased cell migration speed observed in GFP:PaxB-LIMsOnly cells (Figure 6A) where VinculinB localization to cell-substrate adhesions is perturbed (Figures 6F and S6C). These data suggest localization of VinculinB *alongside* PaxillinB specifically dictates the formation of cell-substrate adhesions that inhibit cell migration speed. The recruitment of VinculinB may further recruit other proteins to adhesions, such as actin (Uchida and Yumura, 2004), to enable the proposed molecular “brake” and regulate cell migration speed. Accordingly, in our own data, we observed a population of Paxillin-only punctae (PaxillinB+/VinculinB-) and VinculinB-only punctae (PaxillinB−/VinculinB+) (Figure S6, C-E), and we speculate that these punctae might comprise different subclasses of cell-substrate adhesions with different dynamics and compositions. Unfortunately, due to challenges with low signal to noise and photobleaching, we could not live image mApple-VinculinB at sufficient speeds to quantify lifetime of these different punctae populations. Future research characterizing the dynamics and compositions of these different adhesion classes through improved microscopy resolution and unbiased proteomics approaches, respectively, can shed light on how *Dictyostelium* cells may use different adhesion classes to migrate efficiently.

Our data showing that reducing cell-substrate adhesion number increases cell migration speeds, leads us to hypothesize that while cell-substrate adhesion formation and Paxillin localization to these adhesions is important for cell adhesion across organisms, their role in *promoting* cell migration was likely a Metazoan innovation. Furthermore, these data – plus our data suggesting orthologues of focal adhesion components originated just prior to the divergence of Amoebozoa (Figure 1) – leads us to speculate that the ancestral function of cell-substrate adhesions using these conserved components was primarily for cell adhesion to the underlying substrate. Accordingly, it is worth noting that PaxillinB has additional roles in *Dictyostelium* biology that can shed light on how this speculated innovation of Paxillin and cell-substrate adhesion function might have evolved. Earlier work showed that knockout of PaxillinB impaired *Dictyostelium* multicellular development and spore formation, suggesting that PaxillinB additionally plays a role in cell-cell adhesion and cell differentiation during the transition to aggregate multicellularity (Bukharova et al., 2005; Pribic et al., 2011). In Metazoan systems, cell-cell adhesion and multicellularity are classically thought to be mediated via specialized cell-cell adhesion molecules such as cadherins and catenins, which maintain cell-cell contacts (Gooding et al., 2004; Gul et al., 2017). Catenin molecules are found in non-Metazoan organisms as distant as Amoebozoa and implicated in *Dictyostelium* development, with catenin knockout leading to similar multicellular development defects as PaxillinB knockout (Dickinson et al., 2011; Dickinson et al., 2012). Intriguingly, *Dictyostelium* lacks cadherin molecules, with cadherins appearing to originate just prior to the divergence of Filasterea (Nichols et al., 2012; Suga et al., 2013), the same ancestral branch where FAK appears to have originated (Suga et al., 2013). Taken altogether, these data suggest that *Dictyostelium* PaxillinB may have been a multifunctional protein that was pivotal not only for cell-substrate adhesion formation during cell migration, but also for cell-cell adhesion and development for multicellular aggregation, potentially in conjunction with catenin for the latter functions. Catenins and Paxillin have previously been shown to interact at the interface of adherens junctions and focal adhesions in mammalian endothelial cells through catenin binding with Paxillin LIM domains (Dubrovskyi et al., 2012). To the best of our knowledge, a similar interaction has not been confirmed in *Dictyostelium* but could be indicative of another conserved ancestral function of the Paxillin LIM domain. We further suggest that the subsequent evolution of cadherin and FAK in the last common ancestor to Filasterea may have served as an inflection point at which Paxillin became further specialized for cell-substrate adhesion functions, with more specialization upon the emergence of features such as Paxillin tyrosine phosphoregulation prior to the origin of Metazoa. We believe these specializations, paired with other innovations such as the origin of focal adhesion-specific ECM proteins for specific binding to integrin heterodimers (Hynes, 2012), led to the acquisition of the use of focal adhesions to promote cell migration as has been extensively characterized in mammalian models.

*Dictyostelium*’s combination of a partially conserved focal adhesion toolkit and its amoeboid migration-like features make it an excellent model system to elucidate aspects of cell-substrate adhesion biology that challenge the long-standing framing of focal adhesions and provide a unique window into the early evolution of cell-substrate adhesions structures and their individual molecules. Further characterization of *Dictyostelium* cell-substrate adhesions and their role in regulating cell migration is an exciting avenue of research that will uncover previously unknown forms of cell-substrate adhesion composition, function and regulation in a non-Metazoan model. These findings can then be applied to further inform research on focal adhesions in under-investigated mammalian systems such as non-mesenchymal cells and physiologically relevant contexts. Our work and future investigations are important to re-define the criteria we use for cell-substrate adhesion formation and function during cell migration across various cell types and contexts as well as further understand how this machinery has evolved across Eukaryotes.

## Acknowledgments

We would like to thank all members of the Roh-Johnson lab for discussions and edits during manuscript preparation; Dictybase (RRID:SCR_006643), an NIH-funded Stock Center for providing *Dictyostelium* strains used in this work and for maintaining this valuable community resource; Pascale Charest and the *Dictyostelium* Stock Center for *Dictyostelium* reagents; Nathan Roy and Carole Parent for helpful discussions about *Dictyostelium*; Lillian Fritz-Laylin, Nathan Clark, Nels Elde, and Ellen Leffler for helpful discussion regarding phylogeny and organism choice; and the Cell Imaging Core at the University of Utah for use of Leica Yokogawa CSU-W1 spinning disc confocal microscope, Leica TCS SP8 laser confocal microscope, Nikon Yokogawa CSU-W1 spinning disc confocal microscope, and Nikon Gataca Systems Ilas2 TIRF platform as well as Michael Bridge and Xiang Wang for their assistance.

The work was funded by National Institutes of Health grant R00CA190836 (to M. Roh-Johnson) and National Institute of Allergy and Infectious Diseases grant # 01-00976-5000-59205460 (to BRIDGE UP-HBCU University of Utah), and the National Institute of General Medical Sciences of the National Institutes of Health grant R01GM122917 (M.A. Titus)

## Author contributions

Conceptualization, JCFM, MRJ; Methodology, JCFM, TN, SN, MAT, MRJ; Investigation, JCFM, TN, SN CR, MAT; Formal Analysis, JCFM, TN, CR; Writing, JCFM, MRJ; Funding acquisition, MRJ, BKG, MAT.

## METHODS

### *Dictyostelium* cell culturing

Wild-type AX2 and *paxB*^−^ (DSC Strain DBS0236728) were obtained from the Dictyostelium Stock Center (DSC) and cultured axenically at 21°C in HL5 media (Fey et al., 2007) supplemented with 300 µg/mL Streptomycin Sulfate (Gold Biotechnology) on 10 cm plates. Null cells were selected for with supplementation of 10 µg/mL Blasticidin S (Thermofisher). Transformed cells (see Generation of Transgenic Lines), were selected for with additional supplementation of 20 µg/mL G418 Sulfate (Thermofisher) and/or 50 µg/mL Hygromycin B (Thermofisher).

### Generation of expression plasmids

The wild-type GFP:PaxillinB (UniProt KB accession number Q8MML5) (Nagasaki et al., 2009), RFP:Lifeact (Brzeska et al., 2014), and GFP:TalinB (UniProt KB accession number Q54K81) (Tsujioka et al., 2008) plasmids have been previously described in their respective sources. The authors thank Drs. Masatsune Tsujioka and Taro Q.P. Uyeda (AIST, Tsukuba, Japan) for sharing the GFP:TalinB and GFP:PaxillinB plasmids.

### Cloning of mScarlet:PaxillinB construct

To generate the mScarlet:PaxillinB plasmid, we amplified PaxillinB from the GFP:PaxillinB plasmid for insertion into an mScarlet backbone vector using Gibson assembly. To amplify GFP:PaxillinB (referred to as fragment A1 for Gibson assembly), we used primers A1-PaxillinB-F and A1-PaxillinB-R (Tm = 68°C). To generate mScarlet backbone fragments (referred to as fragments A2 and A3) for Gibson assembly, we used primers A2-mScarlet-Backbone-F and A2-mScarlet-Backbone-R (Tm = 61°C) for fragment A2 and primers A3-mScarlet-Backbone-F and A3-mScarlet-Backbone-R (Tm = 69°C) for fragment A3. PCR fragments were amplified in triplicate using NEB High Fidelity PCR Master Mix with HF Buffer (M0531S), pooled and gel purified using Zymo Gel DNA Recovery Kits (D4002) followed by ethanol precipitation to increase DNA fragment purity. Fragments were Gibson assembled using NEBuilder HiFi DNA Assembly Master Mix (NEB E2621S). Whole plasmid sequencing was performed by Plasmidsaurus using Oxford Nanopore Technology with custom analysis and annotation.

Primer A1-PaxillinB-F:

- 5’-GGTCCAGGTAAGGATCCAATGGCAACAAAAGGATTAAATATG-3’.

Primer A1-PaxillinB-R:

- 5’-TTTAACTAGCTTAAGCAAATAATTTATTATGACAACCTTTACAATATG-3’.

Primer A2-mScarlet-Backbone-F:

- 5’-TTGCTTAAGCTAGTTAAATAAATAAATTATTTAATAAATAAT-3’.

Primer A2-mScarlet-Backbone-R:

- 5’-AATGATGAGCACTTTTAAAGTTCTGCTATG-3’.

Primer A3-mScarlet-Backbone-F:

- 5’-AAAGTGCTCATCATTGGAAAACGTTCTTCG-3’.

Primer A3-mScarlet-Backbone-R:

- 5’-TGGATCCTTACCTGGACCTTTGTATAATTCATCCATACCACC-3’.

### Cloning of truncated PaxillinB constructs

To generate plasmids possessing the truncated PaxillinB molecules outlined in **Figure 4A**, we used the wild-type GFP:PaxillinB as a template to generate backbone and truncated PaxillinB insert fragments for Gibson assembly. For all plasmids, four fragments were amplified for Gibson assembly. For all truncated PaxillinB plasmids, the B3-GFP:PaxB-ALL and B4-GFP:PaxB-ALL backbone fragments were amplified using the B3-GFP-PaxillinB-ALL-F and B3-GFP-PaxillinB-ALL-R primers (Tm = 72°C) for the B3-GFP:PaxB-ALL backbone fragment and the B4-GFP-PaxillinB-ALL-F and 43-GFP-PaxillinB-ALL-R (Tm = 72°C) for the B4-GFP:PaxB-ALL backbone fragment.

For the GFP:PaxB-ΔL4 plasmid, fragments B1-GFP:PaxB-ΔL4 and B2-GFP:PaxB-ΔL4 were amplified using the B1-GFP-PaxillinB-ALL-F and B1-GFP-PaxillinB-ΔL4-R primers (Tm = 72°C) for the B1-GFP:PaxB-ΔL4 fragment and the B2-GFP-PaxillinB-ΔL4-F and B2-GFP-PaxillinB-ALL-R primers (Tm = 72°C) for the B2-GFP:PaxB-ΔL4 fragment.

For the GFP:PaxB-ΔL3 plasmid, fragments B1-GFP:PaxB-ΔL3 and B2-GFP:PaxB-ΔL3 were amplified using the B1-GFP-PaxillinB-ALL-F and B1-GFP-PaxillinB-ΔL3-R primers (Tm = 72°C) for the B1-GFP:PaxB-ΔL3 fragment and the B2-GFP-PaxillinB-ΔL3-F and B2-GFP-PaxillinB-ALL-R primers (Tm = 72°C) for the B2-GFP:PaxB-ΔL3 fragment.

For the GFP:PaxB-ΔL34 plasmid, fragments B1-GFP:PaxB-ΔL34 and B2-GFP:PaxB-ΔL34 were amplified using the B1-GFP-PaxillinB-ALL-F and B1-GFP-PaxillinB-ΔL34-R primers (Tm = 72°C) for the B1-GFP:PaxB-ΔL34 fragment and the B2-GFP-PaxillinB-ΔL34-F and B2-GFP-PaxillinB-ALL-R primers (Tm = 72°C) for the B2-GFP:PaxB-ΔL34 fragment.

For the GFP:PaxB-ΔL234 plasmid, fragments B1-GFP:PaxB-ΔL234 and B2-GFP:PaxB-ΔL234 were amplified using the B1-GFP-PaxillinB-ALL-F and B1-GFP-PaxillinB-ΔL234-R primers (Tm = 72°C) for the B1-GFP:PaxB-ΔL234 fragment and the B2-GFP-PaxillinB-ΔL234-F and B2-GFP-PaxillinB-ALL-R primers (Tm = 72°C) for the B2-GFP:PaxB-ΔL234 fragment.

For the GFP:PaxB-ΔLIMs plasmid, fragments B1-GFP:PaxB-ΔLIMs and B2-GFP:PaxB-ΔLIMs were amplified using the B1-GFP-PaxillinB-ALL-F and B1-GFP-PaxillinB-ΔLIMs-R primers (Tm = 72°C) for the B1-GFP:PaxB-ΔLIMs fragment and the B2-GFP-PaxillinB-ΔLIMs-F and B2-GFP-PaxillinB-ALL-R primers (Tm = 72°C) for the B2-GFP:PaxB-ΔLIMs fragment.

For the GFP:PaxB-LIMsOnly plasmid, fragment B1-GFP:PaxB-LIMsOnly was amplified using the B1-GFP-PaxillinB-LO-F and B1-GFP-PaxillinB-LO-R primers (Tm = 71°C). Fragment B2-GFP:PaxB-LIMsOnly was amplified using the B2-GFP-PaxillinB-LO-F and B2-GFP-PaxillinB-LO-R primers (Tm = 72°C). Fragment B3-GFP:PaxB-LIMsOnly was amplified using the B3-GFP-PaxillinB-LO-F and B3-GFP-PaxillinB-LO-R primers (Tm = 72°C). Fragment B4-GFP:PaxB-LIMsOnly was amplified using the B4-GFP-PaxillinB-LO-F and B4-GFP-PaxillinB-LO-R primers (Tm = 67°C).

PCR fragments were amplified in triplicate using NEB High Fidelity PCR Master Mix with HF Buffer (M0531S), pooled and gel purified using Zymo Gel DNA Recovery Kits (D4002) followed by ethanol precipitation to increase DNA fragment purity. Fragments were Gibson assembled using NEBuilder HiFi DNA Assembly Master Mix (NEB E2621S). Whole plasmid sequencing was performed by Plasmidsaurus using Oxford Nanopore Technology with custom analysis and annotation.

Primer B1-GFP-Paxillin-ALL-F:

- 5’-AATCATGCGAAACGATCCAGCTTGAACATCTTCACCATCC-3’.

Primer B1-GFP-Paxillin-LO-F:

- 5’-ATGGGTAAAGGAGAAGAACTTTTCACTGGAGTTGTCC-3’

Primer B1-GFP-PaxillinB-ΔL4-R:

- 5’-CTCTAGCGAGCTCTTAGCCAGCTTGTTGATGATAATGGACATCACAATAT-3’.

Primer B1-GFP-PaxillinB-ΔL3-R:

- 5’-CAAACTGAAACGGCAAATGTTGAATAGAAATCGGCCTCAC-3’.

Primer B1-GFP-PaxillinB-ΔL34-R:

- 5’-TCTAGCGAGCTCTTAAACGGCAAATGTTGAATAGAAATCGG-3’.

Primer B1-GFP-PaxillinB-ΔL234-R:

- 5’-TCTAGCGAGCTCTTAAAACAATTCTTGATAACATTTTTCAC-3’.

Primer B1-GFP-PaxillinB-ΔLIMs-R:

- 5’-TAGCGAGCTCTTAACGTGATGTTGGTCCTGTTGAATCAATATCT-3’.

Primer B1-GFP-Paxillin-LO-R:

- 5’-ttaAGCAAATAATTTATTATGACAACCTTTACAATATGGTTTACCATTATTAGCG G-3’

Primer B2-GFP-PaxillinB-ΔL4-F:

- 5’-CAAGCTGGCTAAGAGCTCGCTAGAGTCGTCCATCAAT-3’.

Primer B2-GFP-PaxillinB-ΔL3-F:

- 5’-ACATTTGCCGTTTCAGTTTGTTCTGGCTGTGGAAAAG-3’.

Primer B2-GFP-PaxillinB-ΔL34-F:

- 5’-CCGTTTAAGAGCTCGCTAGAGTCGTCCATCAATTG-3’.

Primer B2-GFP-PaxillinB-ΔL234-F:

- 5’-GTTTTAAGAGCTCGCTAGAGTCGTCCATCAATTGTTC-3’.

Primer B2-GFP-PaxillinB-ΔLIMs-F:

- 5’-CATCACGTTAAGAGCTCGCTAGAGTCGTCCATCAATTG-3’.

Primer B2-GFP-Paxillin-LO-F:

- 5’-GGTTGTCATAATAAATTATTTGCTtaagagctcgctagagtcg-3’

Primer B2-GFP-PaxillinB-ALL-R:

- 5’-ATAGTTGCCTGACTCCCCGTCGTGTAGATAACTACGATAC-3’.

Primer B2-GFP-Paxillin-LO-R:

- 5’-tatacaccattggagaggcatgcaccattcct-3’

Primer B3-GFP-PaxillinB-ALL-F:

- 5’-AGTCAGGCAACTATGGATGAACGAAATAGACAGATCGCTG-3’.

Primer B3-GFP-Paxillin-LO-F:

- 5’-caaggaatggtgcatgcctctccaatggtgtatac-3’

Primer B3-GFP-Paxillin-LO-R:

- 5’-ccgttatttgagccatcaagctttggattgtaaattgtt-3’

Primer B3-GFP-PaxillinB-ALL-R:

- 5’-CCGAATCATTGAAACATGGAGGGCAAAGTTTAGAATTAATAACAAC-3’.

Primer B4-GFP-Paxillin-LO-F:

- 5’-atggctcaaataacggtatggaaaagtc-3’

Primer B4-GFP-PaxillinB-ALL-F:

- 5’-GCCCTCCATGTTTCAATGATTCGGTAATCAACAAGAAGTGTG-3’.

Primer B4-GFP-Paxillin-LO-R:

- 5’-TTCTCCTTTACCCATgatccatttttattttttatttaatttaatttatttgttttaag-3’

Primer B4-GFP-PaxillinB-ALL-R:

- 5’-TCAAGCTGGATCGTTTCGCATGATTGAACAAGATGGATTG-3’.

### RNA Extraction and cDNA Library Formation

To extract RNA from *Dictyostelium* cultures, 1e8 *Dictyostelium* cells grown in axenic culture with HL5 were harvested, pelleted at 600 rpm for 5 minutes at 4°C and washed twice with ice-cold KK2 Buffer (161.6 mM KH_2_PO_4_ and 40 mM K_2_HPO_4_, pH = 6.4). The supernatant was removed and cells were then processed for RNA extraction using the QIAgen RNeasy Mini Kit for RNA Purification (QIAgen 74104). RNA purity and concentration was measured using a Nanodrop Spectrophotometer and the RNA was subsequently used to synthesize a whole cDNA library using the Superscript^TM^ IV Vilo^TM^ Master Mix Kit (Invitrogen 11766050).

### Cloning of mApple:VinculinB construct

To amplify the VinculinB (UniProt KB accession number Q54TU2) coding sequence, we used the cDNA library as a template to generate an N-terminal fragment C1 using the C1-VinculinB-F and C1-VinculinB-R primers (Tm = 72°C) and a C-terminal fragment C2 using the C2-VinculinB-F and C2-VinculinB-R primers (Tm = 72°C). To generate mApple backbone fragments (referred to as fragments C2 and C3) for Gibson assembly, we used primers C3-mApple-Backbone-F and C3-mApple-Backbone-R (Tm = 65°C) for fragment C3 and primers C4-mApple-Backbone-F and C4-mApple-Backbone-R (Tm = 70°C) for fragment C4. Fragments C3 and C4 were amplified from an pDM358-mApple (Arthur et al., 2021). PCR fragments were amplified in triplicate using NEB High Fidelity PCR Master Mix with HF Buffer (M0531S), pooled and gel purified using Zymo Gel DNA Recovery Kits (D4002) followed by ethanol precipitation to increase DNA fragment purity. Fragments were Gibson assembled using NEBuilder HiFi DNA Assembly Master Mix (NEB E2621S). Whole plasmid sequencing was performed by Plasmidsaurus using Oxford Nanopore Technology with custom analysis and annotation.

Primer C1-VinculinB-F:

- 5’-ATGGAACAATTATTAGAAGCAATTAGTGATGCAGTATCATCAATGGTATT-3’.

Primer C1-VinculinB-R:

- 5’-AGCCAAGAATAGTTGTTTTGAAAGTTCCTCATTCAATTCTTTG-3’.

Primer C2-VinculinB-F:

- 5’-TTGAATGAGGAACTTTCAAAACAACTATTCTTGGCTAGAATTGC-3’.

Primer C2-VinculinB-R:

- 5’-TTATTTTACTCTAATTTGACCAATTTCTACAGCATTAATAGTACCAACGATAGC-3’.

Primer C3-VinculinB-F:

- 5’-GTCAAATTAGAGTAAAATAAACTAGTTAAATAAATAAATTATTTAATAAATAATAAAAAAACA-3’.

Primer C3-VinculinB-R:

- 5’-ATTTGAAATTACAATGAATGTTTTTCGGTTTTTATTT-3’.

Primer C4-VinculinB-F:

- 5’-AACCGAAAAACATTCATTGTAATTTCAAATGTCGAGG-3’.

Primer C4-VinculinB-R:

- 5’-TTCTAATAATTGTTCCATAGATCTCTTGTACAGCTCGTCCATGC-3’.

### Generation of transgenic lines

*Dictyostelium* cells were transformed via electroporation as previously described (Gaudet et al., 2007). Briefly, axenic cells were cultured in HL5, harvested, pelleted and washed twice with ice-cold H-50 buffer (20 mM HEPES, 50 mM KCl, 1 mM MgSO_4_, 10 mM NaCl, 5 mM NaHCO_3_ and 1 mM NaH_2_PO_4_ • 2H_2_O). Cells were resuspended in H-50 at 1e8 cells/mL and 100 µl of cells were added to a 0.1 cm gap size cuvette with 15 µg of plasmid DNA for single construct transformations or 10 µg of plasmid DNA per construct for double construct transformation. Cells were electroporated via pulsing twice at 0.85 kV, 25 µF with a 0.6 ms τ, with a 5 second pause in between pulses. Cells were rested on ice for 5 minutes post-electroporation before being added to 10 cm plates with fresh HL5 media supplemented with 300 µg/mL Streptomycin Sulfate for 24 hours for recovery. After 24 hours, cells were moved to selection media with 20 µg/mL G418 Sulfate and/or 50 µg/mL Hygromycin B. If null cells were used as a background, 10 µg/mL Blasticidin S was also added to the selection media.

### Preparation of *Dictyostelium* cells for imaging

*Dictyostelium* cells were cultured in HL5 media on 10 cm plates to a confluency of 1-2e6 cells/mL, as measured via Countess II FL (Thermofisher), and a minimum of 10 mLs were harvested. Cells were centrifuged at 600 rpm for 5 minutes at 4°C and resuspended in Development Buffer (DB; 5 mM Na_2_HPO_4_, 5 mM KH_2_PO_4_, 1 mM CaCl_2_, 2 mM MgCl_2,_ pH=6.5) at a concentration of 2.5e^6^ cells/mL. 1e^7^ total cells were plated in a 6 cm plate and allowed to adhere for 30 minutes before being rinsed twice with 2 mL DB and allowed to develop for 3.5 hours. After 3.5 hours, cells were resuspended and 45 µL of cells were added to 955 µL of DB and allowed to adhere in a 35 mm glass bottom dish (FD35-100, World Precision Instruments) for 30 minutes before moving on to imaging.

### Dictyostelium imaging

Spinning disc confocal fluorescence microscopy was done using either a Leica HC PL APO 63×/1.40 oil immersion objective with 2× zoom on a Leica Yokogawa CSU-W1 spinning disc confocal scanner unit with iXon Life 888 EMCCD camera and VisiView software (RRID:SCR_022546) for spinning disc confocal microscopy **(for images and data generated for Figures 2B, 2E, 6A and 6B)** or a Nikon PLAN APO λD 60X/1.42 oil immersion objective with 1X zoom on a Nikon Ti2 microscope with a Yokogawa CSU-W1 spinning disc confocal scanner unit, Gataca Systems Live-SR super resolution module and Kinetix 22 Scientific CMOS (sCMOS) camera for super resolution spinning disc confocal microscopy with NIS AR software (RRID:SCR_014329) **(for images and data generated for Figures 2D, 2G, 2H, 5B, 5C, 6F, 6G, S1B-D, S3A-D, S4, S5 and S6)**. Laser scanning confocal microscopy was done using Leica HC PL APO CS2 63×/1.40 oil immersion objective with 2X zoom on a Leica TCS SP8 laser confocal microscope with photomultiplier tubes (PMT) detectors and Leica Application Suite X software (RRID:SCR_013673) **(for images generated for Figure 2C).** TIRF microscopy was done using a Nikon 60X Apochromat TIRF oil objective with 1X zoom on a Nikon Ti2 microscope with Gataca Systems Ilas2 TIRF platform and iXon Ultra 897 EMCCD camera and NIS AR software **(for images generated for Figure S1)**. For all microscopy, *Dictyostelium* cells were in development buffer (DB).

### LatrunculinA spike-in

*Dictyostelium* cells undergoing random cell migration were imaged in a single plane every 5 sec for 10 min. 1 ml of 2 µM LatrunculinA (Sigma Aldrich) was spiked-in to the imaging dish at the 4 min mark to bring the final concentration of LatrunculinA to 1 µM. Imaging was continued for the remaining 6 min to observe effect of spike-in.

### *Dictyostelium* cell spreading

Wild-type and *paxB^−^ Dictyostelium* cells were captured using Differential Interference Contrast (DIC) microscopy (**for images and data generated for Figures 3A, B)**. Image files were blinded using ImageJ’s (RRID:SCR_003070) (Schindelin et al., 2012) Blind Analysis Tools and processed using the Trainable Weka Segmentation plug-in (Arganda-Carreras et al., 2017) on ImageJ to classify cells and create whole cell binary map. Cell area was subsequently measured using the Measurement tool on ImageJ.

### Creation and priming of microfluidics chamber

The microfluidic device was designed using AutoCAD (RRID:SCR_021072) and was fabricated with laser-cut laminates. The idea behind the device design is to have a widening channel where the fluid flow has a range of speeds depending on the location along the channel. The narrower end has a faster speed and exposes the cells to higher shear stress, while the wider end results in a slower fluid speed, which, in turn, exposes the cells to a lower shear stress. This helps cover a range of shear stress on a single run. The channel width at the narrowest end was designed to be 1.6 mm and at the widest end was set to be 18.35 mm. The channel was widened at a constant rate along the total channel length of 64 mm. Multiple pillars were also designed along the channel to prevent the channel roof from collapsing during the run and to act as markers for image locations. The device bottom, channel cutout and the channel roof were laser-cut from a standard Petri dish, 3M 9965 and 3M 9960, respectively. 3M 9965 is a white biocompatible polyester carrier with hydrophobic adhesives on both sides, while 3M 9960 is a transparent biocompatible polyester carrier with no adhesives on either side; 3M 9965 and 3M 9960 were bonded first. To make the petri dish surface hydrophilic, it was plasma treated before bonding it to the other side of the 3M 9965. Finally, one inlet and three outlet PDMS ports were attached to the openings on the narrowest and widest ends of the 3M 9960 channel roof, respectively. The device was then placed in an oven at a temperature of 70°C for 10 minutes.

During the experiment, a pipette tip was inserted into the inlet PDMS port to act as an inlet reservoir and 100 µl of DB was loaded into the reservoir to prime the channel. Keeping the outlet ports open, the DB was allowed to naturally flow into the device to get rid of the air trapped inside. Next, three tubes were inserted into the outlet ports. And three primed 5 ml syringes were connected to the tubes. The syringes were then placed into a multichannel syringe pump (KD Scientific KDS220) to control the flow rates at which the sample in the inlet reservoir would be pulled into and out of the device.

### *Dictyostelium* microfluidics assay and detachment quantification

*Dictyostelium* cells were prepared as previously stated in “Preparation of *Dictyostelium* cells for imaging” and 150,000 cells in 100ul of DB were loaded into the pipette tip located at the loading reservoir. Once loaded, a flow rate of 50 µl/min for 2 minutes was used to gently pull the cells into the chamber and fill it with fluid. An extra 100 µl of DB was added when the initial 100 µl was almost depleted to avoid introducing air bubbles into the chamber. Cells were allowed to adhere for 30 minutes before applying an initial flow rate of 250 µl/min for 1 min to remove any remaining cells that did not adhere. Cells were then consecutively subjected to flow rates of 500 µl/min, 750 µl/min, 1000 µl/min, 1250 µl/min and 1500 µl/min for 2 minutes at each rate to test adhesion. During the microfluidics assay, a pre-chosen field of view was imaged throughout the assay every two seconds using an 100X objective on an AmScope IN50 Inverted Compound Microscope with AmScope FMA050 MU1803 Camera and Software. To calculate percent detachment, the initial number of cells after the initial flow rate of 250 µl/min was counted using the ImageJ Multi-Point tool. The number of cells left at the end of each 2 minute flow rate interval was then counted and subtracted from the initial number of cells to calculate the percent of cells that had detached by the end of each flow rate interval.

### *Dictyostelium* chemotaxis assay

Chemotactically competent Dictyostelium cells were prepared by harvesting 1 x 10^8^ cells and washing them twice with DB (450 x g for 3 min). Cells were resuspended to a final density of 2 x 10^7^ cells/mL, shaken on a rotating shaker for 1 hr at 200 rpm, then pulsed with 100nM cAMP every 6 min for 3 hr. Cells were collected and washed with DB prior to assay. Chemotaxis was assayed using µ-Slide Chemotaxis chamber (IBIDI, Martinsried, Germany). A suspension of 1 x 10^6^ cells/mL chemotactically competent cells was seeded into one of the two reservoirs and allowed to adhere for 10 min. The other reservoir was filled with a 1 µM cAMP solution that included 2 µg/mL final of Alexa fluor 597 hydrazide (Invitrogen). The gradient was allowed to establish for 20 minutes before acquisition. Imaging was done with a Nikon Ti2-e equipped with Hamamatsu ORCA-Fusion BT CMOS camera (C15440-20UP) and CrestOptics X-Light V3 Spinning disc unit and Nikon Plan Fluor 40x/1.30 oil-immersion objective. Images were captured using NIS-Element AR software (v5.42.04). Cells migrating through the observatory channel were imaged at 15 sec intervals for a total of 15 min in DIC and 488 nm channels. The cAMP gradient was captured in the 561 nm channel at the beginning of each acquisition and validated by performing a horizontal line scan analysis for pixel intensity across the field of view in ImageJ.

### Identification and quantification of PaxillinB punctae number and size for punctae analysis and co-localization

PaxillinB punctae were identified and quantified using a custom-made ImageJ processing pipeline. Briefly, fluorescent images of *Dictyostelium* cells expressing wild-type or truncated GFP-tagged PaxillinB were auto-thresholded and background noise was removed through ImageJ’s NaN Background tool. After background removal, auto-threshold was performed again and the mean fluorescence intensity (MFI) and standard deviation (SD) was then measured for each cell. To identify and quantify punctae, thresholding was used with a value of MFI + 3 SD used as the lower limit and 65535 used as the upper limit. For cells expressing the PaxillinB-LIMsOnly construct, we instead used a thresholding value of MFI + 2 SD due to a high signal-to-noise ratio as a result of the increased localization to ventral surface punctae. Punctae identified via thresholding were then analyzed using the Analyze Particles command, filtering to keep punctae between 0.02 µm^2^ and 0.5 µm^2^. Total number of punctae per cell was displayed in the Summary table while individual punctae were displayed in the Results table after the pipeline was executed. The full ImageJ macro for punctae identification and quantification is available on GitHub (https://github.com/rohjohnson-lab/Fierro_Morales_et_al_2024).

### Identification of VinculinB punctae and quantification of punctae number and co-localization

VinculinB punctae were identified and quantified using the ImageJ processing pipeline, with a thresholding value of thresholding value of MFI + 2 SD used due to a low signal-to-noise ratio for VinculinB. To calculate co-localization between VinculinB and PaxillinB punctae in the same cell, the ROIs of identified PaxillinB and VinculinB punctae in their respective channel were manually analyzed for overlap to identify and quantify both co-localizing and individually localizing punctae in each channel.

### *Dictyostelium* random cell migration analysis

*Dictyostelium* cells undergoing random cell migration were imaged in a single plane every 5 sec for 5 min in the GFP channel. Migration was tracked using auto thresholding and binary tracking of whole cells in NIS-Elements AR Advanced 2D Tracking software. Final speeds were calculated by averaging the speeds between each time point of the time-lapse video. Mean square displacement was calculated using the DiPer computer program (Gorelik and Gautreau, 2014).

### *Dictyostelium* chemotactic migration analysis

Directed cell migration was analyzed by first generating cell masks from the 488 channel by thresholding in FIJI and then tracking was done via TrackMate v7.13.2 (Ershov et al., 2022). The TrackMate simple LAP tracker was used with 25.0 microns linking max distance, 10.0 microns gap-closing max distance, and 5 gap-closing frame gap. Spider plots were generated from TrackMate _spots and _tracks outputs via GitHub CellTracksColab TrackMate repository. Only datasets with 80% of PaxB WT rescued cells migrating toward the cAMP gradient as seen in the spider plots were included in the speed analysis. Statistical analysis was done in Prism 10 (GraphPad) using mean track speed from TrackMate _tracks outputs.

### PaxillinB punctae lifetime analysis

On NIS-Elements AR, a “Regional Maxima” 3 x 3 kernel operation (Sysko and Davis, 2010) was applied to timelapse videos of individual *Dictyostelium* cells undergoing random cell migration imaged every 750 milliseconds to remove background noise and mitigate thresholding issues due to uneven illumination. Auto-thresholding was applied to the whole video, as well as edge smoothing and clean-up to remove single pixels, to define PaxillinB punctae as distinct binary objects. NIS-Elements AR Advanced 2D Tracking was then used to track PaxillinB binaries and mean fluorescent intensities (MFI) across each frame. For Advanced 2D Tracking, punctae present in the first or last frame of the video were removed and a minimum duration of eight frames (6 seconds) was required. Lastly, manual curation of identified punctae was performed to remove any punctae binaries that merged or skipped frames. To quantify lifetime using generated PaxillinB binaries data, for each binary the MFI value from the first frame was subtracted from all other frames. A five-frame running average was calculated to reduce noise and improve curve fitting for lifetime curve generation on Excel using methods previously described (Stehbens and Wittmann, 2014). Briefly, data points were placed on Excel and the “Solver” add-in was used to generate adhesion assembly and disassembly curves based on fit to logistic function and exponential decay, respectively. A quantitative lifetime was then calculated based on the t-half of both the assembly and disassembly fit.

### Bidirectional homology queries

For all bidirectional homology queries, we used the *Homo sapiens* α*-*Integrin, β*-*Integrin, Vinculin, Talin, Zyxin, Paxillin, α*-*Actinin, c-SRC, and FAK protein sequences obtained from UniProt (RRID:SCR_002380) as initial queries. First, sequence homology was done using the Protein basic local alignment search tool (BLASTp) (RRID:SCR_001010) to blast the *H.sapiens* protein sequences against genomes of candidate organisms in the National Center for Biotechnology Information (NCBI) (SCR_002474), Joint Genome Institute (JGI) (RRID:SCR_002383) and Ensembl (RRID:SCR_002344) genome databases (Altschul et al., 1990; Martin et al., 2023; Nordberg et al., 2014), using an e-value threshold of 10^−10^. Reciprocal searches were performed using identified putative orthologue sequences from candidate organisms against the *H.sapiens* genome. For domain family-based analyses, we input the *H.sapiens* protein sequences into the HMMER (RRID:SCR_005305) webserver (Potter et al., 2018) using the built-in pHMMER and HMMSCAN tools to identify putative orthologues in candidate organisms and confirm domain architecture of input queries, respectively. This process was repeated using identified putative orthologues from candidate species to test for reciprocity to the *H.sapiens* genome and confirm predicted domain architecture of candidate orthologue queries (see Table 1 for UniProtKB amino acid sequence accession numbers used for both BLASTp and pHMMER analyses). For structural homology, predicted Alphafold (RRID:SCR_023662) structures (Jumper et al., 2021; Varadi et al., 2022) of the *H.sapiens* adhesion molecules were used and structural homologs were searched for in candidate organisms using the Foldseek (van Kempen et al., 2023) (RRID:SCR_018390) webserver with the AlphafoldDB/UniProt50 v4, AlphafoldDB/Swiss-Prot v4, AlphafoldDB/Proteome v4, CATH50 4.3.0 (RRID:SCR_007583), Protein Data Bank (PDB) 100 20231120 (SCR_012820), MGnify-ESM30 v1 (RRID:SCR_016429), and GMGCL 2204 databases (Berman et al., 2000; Coelho et al., 2022; Knudsen and Wiuf, 2010; Richardson et al., 2023; Varadi et al., 2022) and 3Di/AA structural alignment methods. Reciprocal searches were performed using identified putative orthologue structures from candidate organisms as the input query against *H.sapiens* proteosome (see Table 2 for predicted AlphaFoldDB structure accession numbers used for Foldseek analysis). If no homologs were detected using the *H.sapiens* sequences or structures for a candidate organisms, queries were repeated using previously identified putative orthologues from other organisms more closely related to the candidate organisms in order to account for distant homology.

### Homology searches for LIM domain possessing proteins

Methods described in the “Bidirectional homology queries” section were repeated, this time using the *Homo sapiens* sequences and structures for Paxillin, Leupaxin, LIMS1, LIMS2, PDLIM7, Testin and Zyxin as initial queries (see Table 3 for UniProtKB amino acid sequence accession numbers and AlphaFoldDB structure accession numbers used).

### Multiple sequence alignments and phylogenetic analysis

Protein sequence were aligned using the MAFFT (Katoh and Standley, 2013)(RRID:SCR_011811) plug-in on Geneious Prime software (Dotmatics) (RRID:SCR_010519). A maximum likelihood (ML) phylogenetic tree was then inferred from the resulting multiple sequence alignment using PHYML (Guindon et al., 2010) (RRID:SCR_014629) plug-in with an LG substitution model and 100 bootstraps replicates. Trees were visualized using FigTree v1.4.4 software (http://tree.bio.ed.ac.uk/software/figtree/) (RRID:SCR_008515)

### Paxillin predicted structural analysis

PDB files for the predicted Alphafold structures of *H.sapiens* Paxillin (AlphafoldDB accession number AF-P49023-F1-v4) and *D.discoideum* PaxillinB (AlphafoldDB accession number AF-Q8MML5-F1-v4) molecules were obtained from the Alphafold Protein Structure Database (Varadi et al., 2022). The PDB files were visualized in USCF ChimeraX software developed by the Resource for Biocomputing, Visualization, and Informatics at the University of California, San Francisco, with support from National Institutes of Health R01-GM129325 and the Office of Cyber Infrastructure and Computational Biology, National Institute of Allergy and Infectious Diseases (Meng et al., 2023). Structures were aligned using the Matchmaker feature and a Smith-Waterman alignment algorithm.

### Graphical representation and statistical analysis

All graphs were generated from GraphPad Prism (v10) (RRID:SCR_002798) and Microsoft Excel (v16.43) (RRID:SCR_016137). Statistical and normality analyses were performed using Prism (v8, GraphPad). D’Agostino-Pearson, Shapiro-Wilk, Anderson-Darling and Kolmogorov-Smirnov normality tests were run prior to statistical analyses to test for normality of datasets. Outliers were identified and removed using Prism’s “Identify Outliers” tool using the ROUT method with a coefficient Q value of 1% (Motulsky and Brown, 2006) for **Figures 2B, 2H (punctae number), 3B, 5A (except for columns PaxB-ΔL234 and PaxB-ΔLIMs) 5B, 5C, 6E, 6G, S1C, and S5, S6A, S6D (only for Pax+/VinB− and Pax+/VinB+ columns for PaxB, PaxB-ΔL4, PaxB-ΔL34 and the Pax+/VinB− column of PaxB-LIMsOnly), and 6E (except for columns GFP and PaxB-ΔLIMs).** For other graphs, outliers were not calculated because we had already defined criteria during pipeline analysis **(Figures 2H area, S1D, S3A, S6B),** or there was a large number of zero values **(>50%; Figure 5A columns PaxB-ΔL234 and GFP:PaxB-ΔLIMs, Figure 6G columns PaxB-ΔLIMs and PaxB-LIMsOnly, Figure S6D (all PaxB-−/VinB+ columns, PaxB+/VinB− columns of GFP and PaxB-ΔLIMs and Pax+/VinB+ columns of GFP, PaxB-ΔLIMs, and PaxB-LIMsOnly) and Figure S6E (Columns GFP and PaxB-LIMsOnly.** N-values of biological replicates, the number of cells, and specific statistical tests used are found in the figure legends. Only comparisons that are statistically significant are shown on graphs. Correlation analyses were performed by running the simple linear regression analysis function in Prism.

