## Supplemental Files for "Reduced cell-substrate adhesion formation promotes cell migration in *Dictyostelium*"

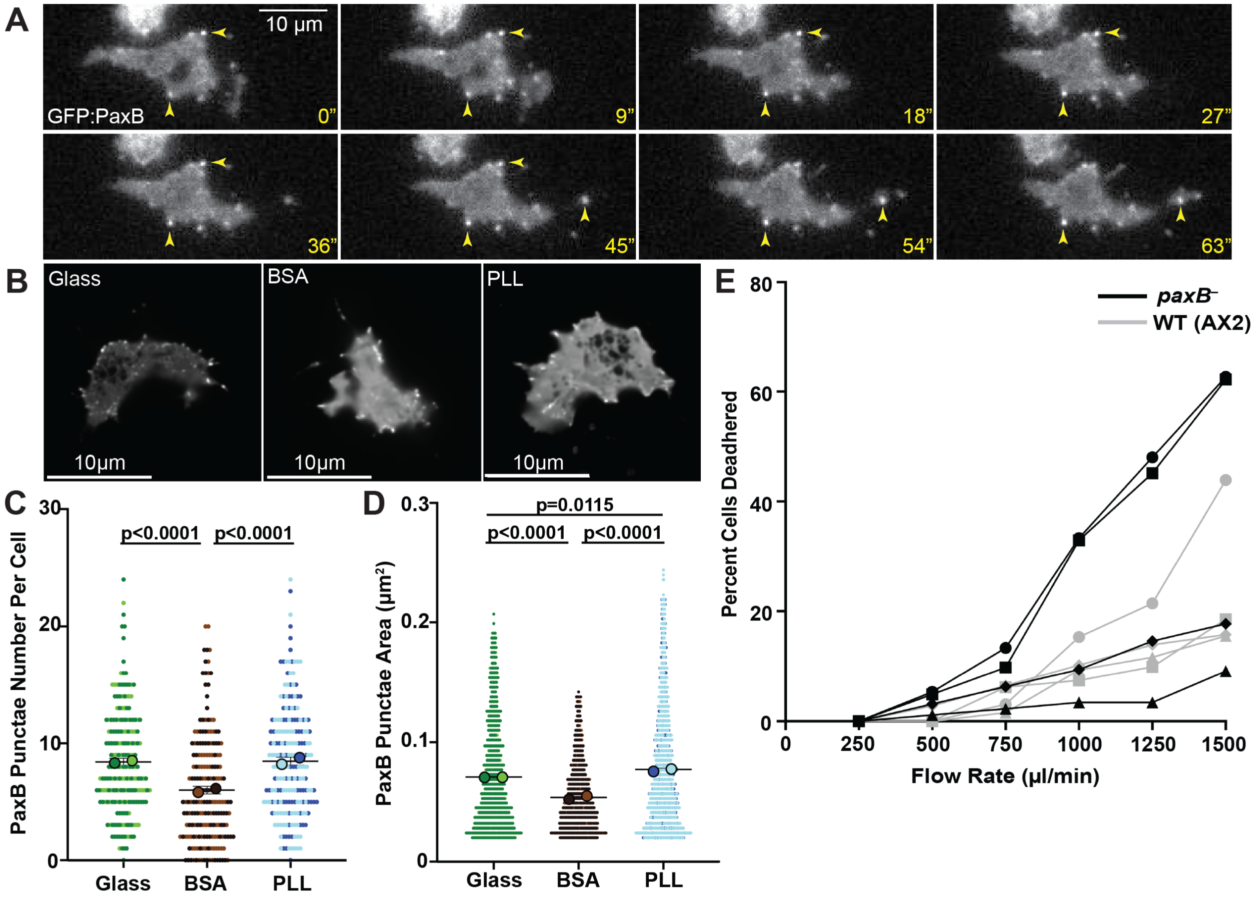


**Supplemental Figure 1:** PaxillinB punctae formation and size at the ventral surface are affected by the underlying substrate, and *paxB*⁻ cells exhibit reduced cell-substrate adhesion.

(A) Representative timelapse total internal reflection fluorescence (TIRF) microscopy images of *paxB*⁻/act15/GFP:PaxB;RFP:Lifeact *Dictyostelium* cells. Yellow arrowheads point to PaxillinB punctae forming at the cell ventral surface during cell migration. Time indicated in seconds. (B) Representative fluorescent confocal microscopy images of *paxB*⁻/act15/GFP:PaxB;RFP:Lifeact *Dictyostelium* cells plated on glass, bovine serum albumin (BSA) or poly-L-lysine (PLL). (C) Quantification of number of PaxillinB punctae per cell in *Dictyostelium* cells plated on indicated surfaces. n = 201 (Glass), 197 (BSA), and 204 (PLL) cells across n=2 biological replicates per cell line. Mean +/- SEM; Kruskal-Wallis Test. (D) Quantification of PaxillinB individual punctae area in *paxB*⁻/act15/GFP:PaxB *Dictyostelium* cells plated on indicated surfaces. n = 1705 (Glass), 1207 (BSA), and 1745 (PLL) punctae across n=2 biological replicates per cell line. Mean +/- SEM, Kruskal-Wallis Test. (E) Individual replicates used for quantification of cell detachment of *paxB*⁻ and wildtype (AX2) *Dictyostelium* cells to compare cell adhesion capability as seen in Figure 3D. Different shapes correspond to different paired replicates. For all graphs, only comparisons that are statistically significant are shown.

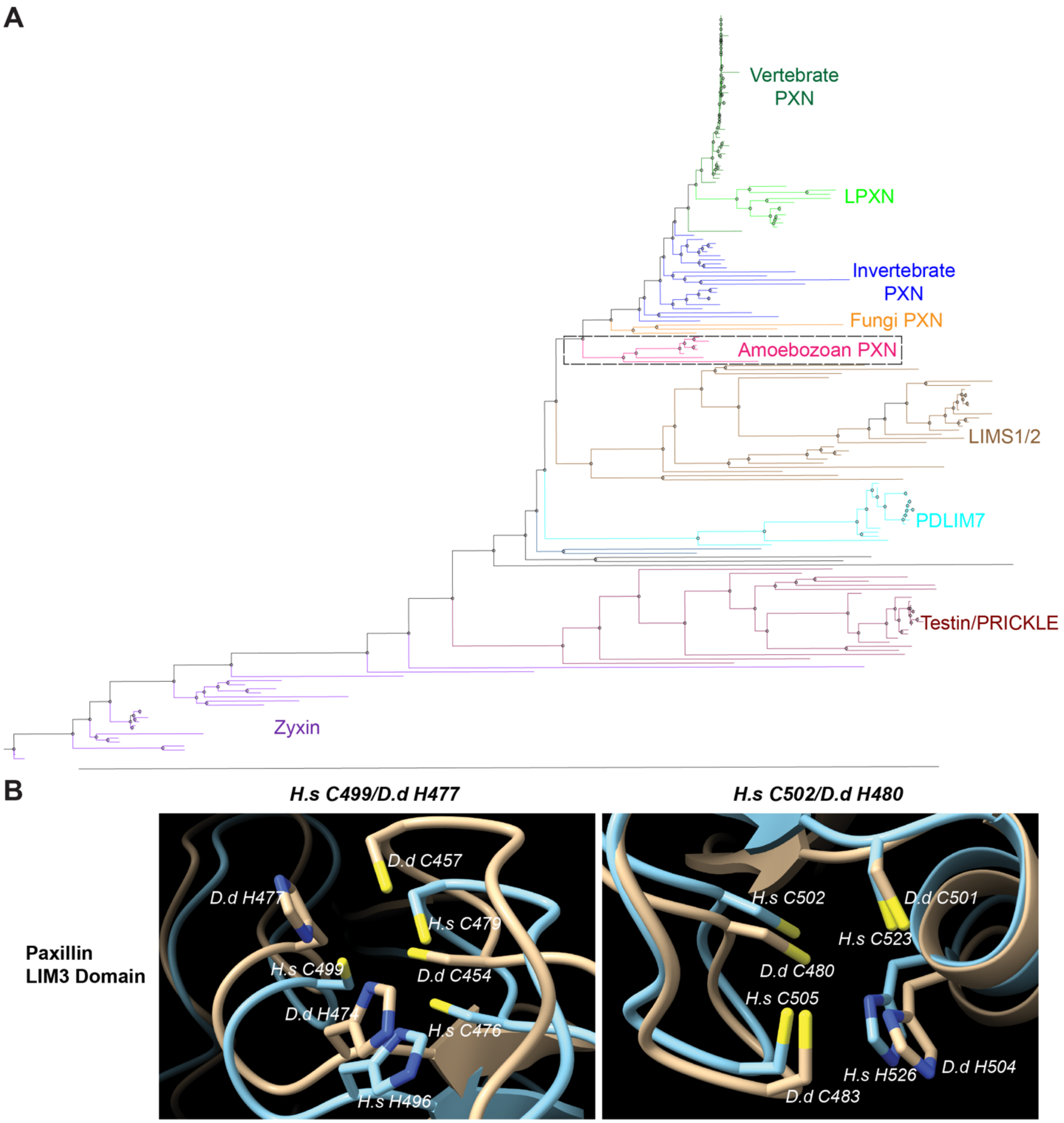


**Supplemental Figure 2:** *Dictyostelium* Paxillin LIM domains cluster with LIM domains of known Paxillin family members and possess conserved zinc binding residues for adhesion localization. (A) PhyML 3.0 was used to generate a phylogenetic tree of LIM domains from proteins possessing multiple LIM domains across Eukaryotic species. The different proteins (Paxillin, Leupaxin, LIMS1/2, PDLIM7, Testin/Prickle and Zyxin) are labeled and color-coded to the right of the tree. For Paxillin molecules, we further highlighted the different phyla. The topology indicates that LIM domains from putative Amoebozoan Paxillin molecules (highlighted by black dashed box) cluster more closely with LIM domains from other Paxillin and Paxillin family members, such as Leupaxin (LPXN), and not with LIM domains from other LIM domain possessing proteins (such as LIMS1/2, PDLIM7, Testin/Prickle and Zyxin). (B) Alignment of Alphafold-predicted structures of the LIM3 domains of *Dictyostelium* PaxillinB (brown) and Human Paxillin (light blue) show conservation of two different zinc-coordinating pockets within LIM3 – Human C499/*Dictyostelium* H477 and Human C502/*Dictyostelium* H480. The zinc-coordinating pockets are formed by the cysteine (highlighted in yellow) and histidine (highlighted in dark blue) residues. Predicted structures were visualized and aligned using ChimeraX Matchmaker.

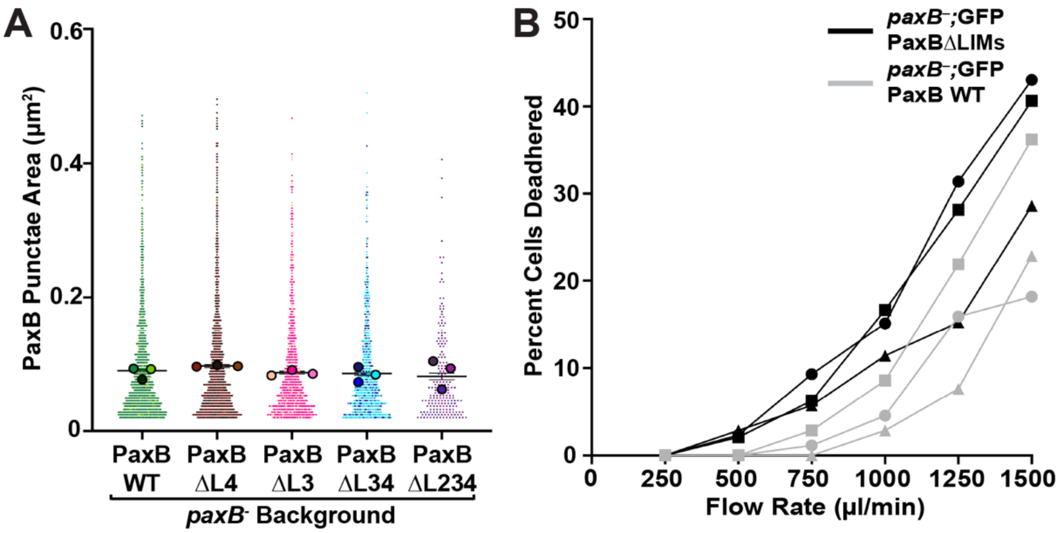


**Supplemental Figure 3:** Perturbation of PaxillinB LIM domains decreases cell adhesion but does not affect PaxillinB-positive adhesion size.

(A) Quantification of PaxillinB punctae area of individual punctae in *paxB⁻ Dictyostelium* cells overexpressing PaxillinB molecules shown in Figure 5A. n = 1611 (PaxB-WT), 1439 (PaxB-ΔL4), 1112 (PaxB-ΔL3), 807 (PaxB-ΔL34), and 199 (PaxB-ΔL234) PaxillinB punctae across n=3 biological replicates per cell line. Mean +/- SEM, Kruskal-Wallis Test. Statistical analyses showed no significant differences between cell lines. (B) Individual replicates used for quantification of cell detachment of *paxB⁻ Dictyostelium* cells overexpressing either GFP:PaxB-WT or GFP:PaxB-ΔLIMs proteins to compare cell adhesion capability as seen in Figure 5D. Different shapes indicate different paired replicates.

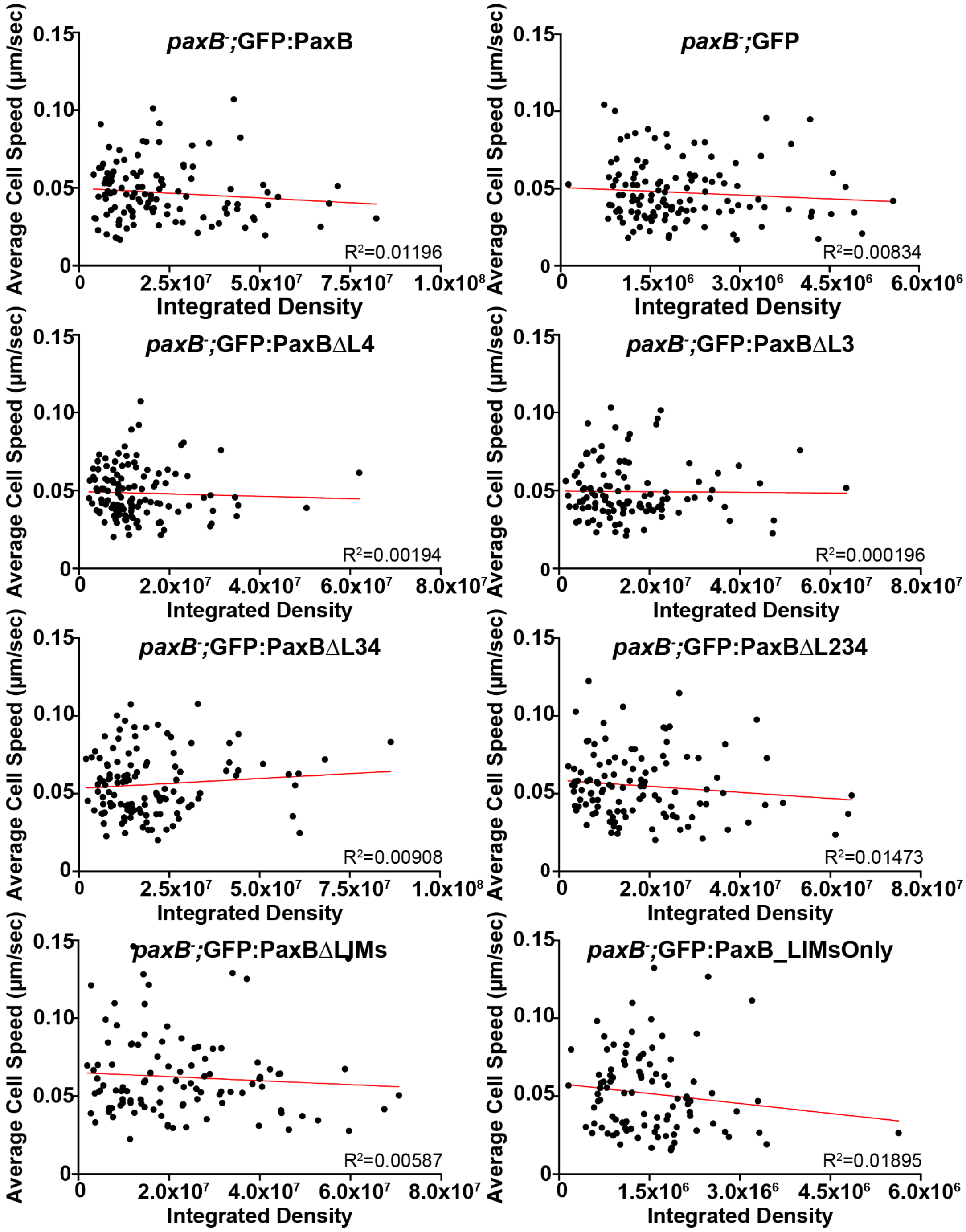


**Supplemental Figure 4:** *Dictyostelium* cell migration speed is not influenced by PaxillinB expression level.

Scatterplot graphs comparing cell migration speed to PaxillinB expression level of individual *Dictyostelium* cells graphed in Figure 6A. GFP:PaxillinB expression was measured by quantifying total integrated GFP intensity of each cell. Correlation analyses of cell migration speed and integrated density measurements of individual cells suggest there is no correlation between PaxillinB expression level and cell migration speed for any of the *Dictyostelium* cell lines.

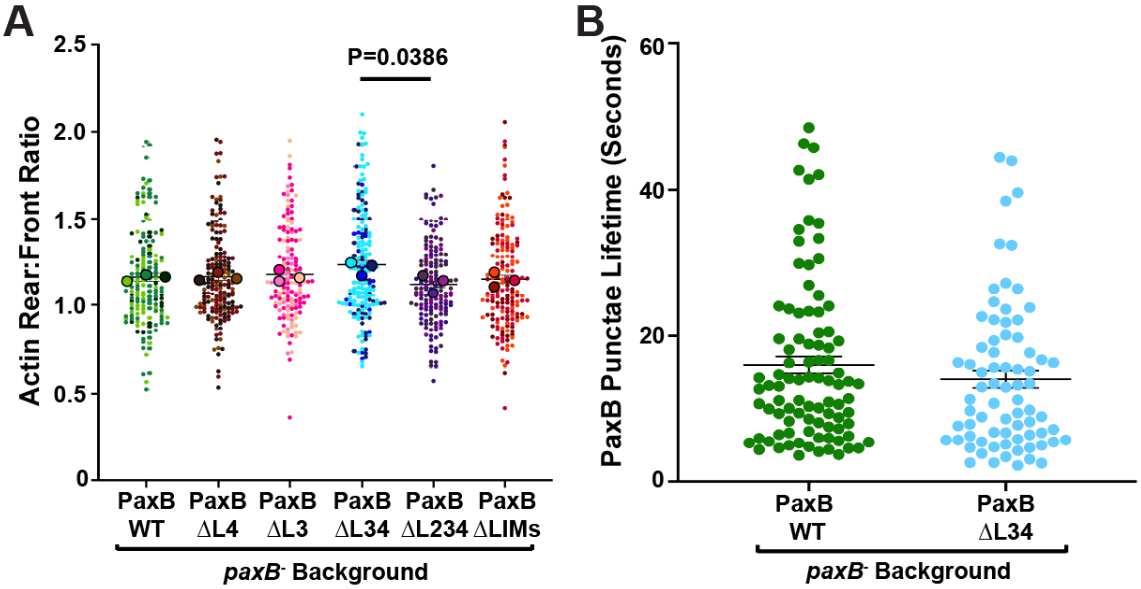


**Supplemental figure 5:** Perturbation of PaxillinB localization does not trigger a transition to amoeboid-based motility or significantly affect adhesion dynamics during random migration. (A) Quantification of the ratio of actin, using the Mean Fluorescence Intensity (MFI) of RFP-tagged Lifeact, at the rear versus the front of *paxB*$^{-}$ *Dictyostelium* cells overexpressing PaxillinB molecules shown in Figure 5A. n = 158 (PaxB-WT), 175 (PaxB-ΔL4), 159 (PaxB-ΔL3), 188 (PaxB-ΔL34), 161 (PaxB-ΔL234), and 174 (PaxB-ΔLIMs) cells across n=3 biological replicates per cell line. Mean +/- SEM. Kruskal-Wallis Test. (B) Quantification of PaxillinB punctae lifetime during timelapse imaging of *paxB*$^{-}$ *Dictyostelium* cells overexpressing wildtype PaxillinB (PaxB-WT) or truncated PaxillinB missing the LIM3 and LIM4 domains (PaxB-ΔL34). n = 93 (PaxB-WT) and 79 (PaxB-ΔL34) punctae across n=1 biological replicates per cell line. Student t-test. Only comparisons that are statistically significant are shown on the graphs.

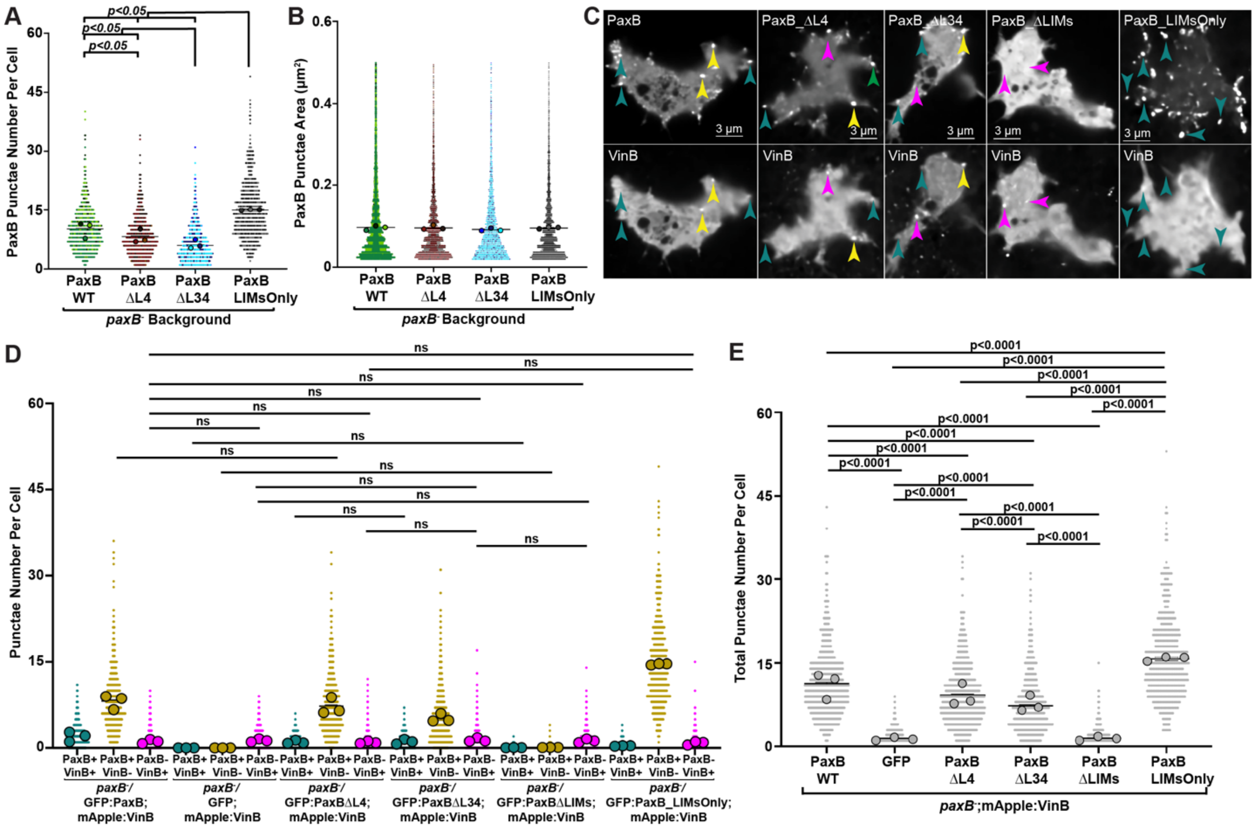


**Supplemental Figure 6:** *Dictyostelium* cells expressing PaxillinB LIM domains alone have increased PaxillinB punctae but no co-localization with VinculinB. (A) Quantification of the number of PaxillinB punctae in *paxB*$^{-}$ *Dictyostelium* cells overexpressing VinculinB and PaxillinB molecules shown in Figure 6, F and G. n = 745 (PaxB-WT), 646 (PaxB-ΔL4), 676 (PaxB-ΔL3), and 552 (PaxB-LIMsonly) cells across n=3 biological replicates per cell line. Mean +/- SEM, Kruskal-Wallis Test. (B) Quantification of the area of individual PaxillinB punctae in *paxB*$^{-}$ *Dictyostelium* cells overexpressing VinculinB and PaxillinB molecules shown in Figure 6F. n = 6052 (PaxB-WT), 4681 (PaxB-ΔL4), 3390 (PaxB-ΔL3), and 8068 (PaxB-LIMsOnly) PaxillinB punctae across n=3 biological replicates per cell line. Mean +/- SEM, Kruskal-Wallis Test. Statistical analyses showed no significant difference between cell lines. (C) Representative images of timelapse fluorescent confocal microscopy of *paxB*$^{-}$ *Dictyostelium* cells overexpressing mApple-VinculinB and the PaxillinB molecules indicated in each image. Golden arrowheads point to PaxillinB-positive and VinculinB-positive punctae (PaxB+;VinB+). Teal arrowheads point to PaxillinB-positive and VinculinB-negative punctae (PaxB+;VinB-). Magenta arrowheads point to PaxillinB-negative and VinculinB-positive punctae (PaxB-;VinB+). (D) Quantification of PaxB+;VinB+, PaxB+;VinB-, or PaxB-;VinB+ punctae per cell in (C). Due to space constrictions, only non-significant differences are shown. All other comparisons are statistically significant ($p<0.05$). (E) Quantification of the total number of punctae per cell in (C). Only comparisons that are statistically significant are shown. For both (D and E) n = 745 (PaxB-WT), 646 (PaxB-ΔL4), 676 (PaxB-ΔL3), 593 (PaxB-ΔLIMs), 552 (PaxB-LIMsOnly), and 620 (Empty) cells across n=3 biological replicates per cell line. Mean +/- SEM.

**<u>Supplemental Tables for Accession Numbers</u>**

**Table 1**

| Protein Name | Organism | Database | Accession Number |
|---|---|---|---|
| Paxillin (PXN) | *Homo sapiens* | UniProtKB | P49023 |
| Vinculin (VCL) | *Homo sapiens* | UniProtKB | P18206 |
| Talin-1 (TLN1) | *Homo sapiens* | UniProtKB | Q9Y490 |
| Talin-2 (TLN2) | *Homo sapiens* | UniProtKB | Q9Y4G6 |
| Zyxin (ZYX) | *Homo sapiens* | UniProtKB | Q15942 |
| Focal Adhesion Kinase 1 (FAK1) | *Homo sapiens* | UniProtKB | Q05397 |
| Focal Adhesion Kinase 2 (FAK2) | *Homo sapiens* | UniProtKB | Q14289 |
| Proto-oncogene tyrosine-protein kinase SRC (c-SRC) | *Homo sapiens* | UniProtKB | P12931 |
| Integrin Alpha 1 (ITGA1) | *Homo sapiens* | UniProtKB | P56199 |
| Integrin Alpha 2 (ITGA2) | *Homo sapiens* | UniProtKB | P17301 |
| Integrin Alpha 3 (ITGA3) | *Homo sapiens* | UniProtKB | P26006 |
| Integrin Alpha 4 (ITGA4) | *Homo sapiens* | UniProtKB | P13612 |
| Integrin Alpha 5 (ITGA5) | *Homo sapiens* | UniProtKB | P08648 |
| Integrin Alpha 6 (ITGA6) | *Homo sapiens* | UniProtKB | P23229 |
| Integrin Alpha 7 (ITGA7) | *Homo sapiens* | UniProtKB | Q13683 |
| Integrin Alpha 8 (ITGA8) | *Homo sapiens* | UniProtKB | P53708 |
| Integrin Alpha 9 (ITGA9) | *Homo sapiens* | UniProtKB | Q13797 |
| Integrin Alpha 10 (ITGA10) | *Homo sapiens* | UniProtKB | O75578 |
| Integrin Alpha 11 (ITGA11) | *Homo sapiens* | UniProtKB | Q9UKX5 |

| | | | |
|---|---|---|---|
| Integrin Alpha D (ITGAD) | *Homo sapiens* | UniProtKB | Q13349 |
| Integrin Alpha E (ITGAE) | *Homo sapiens* | UniProtKB | P38570 |
| Integrin Alpha L (ITGAL) | *Homo sapiens* | UniProtKB | P20701 |
| Integrin Alpha M (ITGAM) | *Homo sapiens* | UniProtKB | P11215 |
| Integrin Alpha V (ITGAV) | *Homo sapiens* | UniProtKB | P06756 |
| Integrin Alpha-IIb (ITGA2B) | *Homo sapiens* | UniProtKB | P08514 |
| Integrin Alpha X (CD11c) | *Homo sapiens* | UniProtKB | P20702 |
| Integrin Beta 1 (ITGB1) | *Homo sapiens* | UniProtKB | P05556 |
| Integrin Beta 2 (ITGB2) | *Homo sapiens* | UniProtKB | P05107 |
| Integrin Beta 3 (ITGB3) | *Homo sapiens* | UniProtKB | P05106 |
| Integrin Beta 4 (ITGB4) | *Homo sapiens* | UniProtKB | P16144 |
| Integrin Beta 5 (ITGB5) | *Homo sapiens* | UniProtKB | P18084 |
| Integrin Beta 6 (ITGB6) | *Homo sapiens* | UniProtKB | P18564 |
| Integrin Beta 7 (ITGB7) | *Homo sapiens* | UniProtKB | P26010 |
| IIntegrin Beta 8 (ITGB8) | *Homo sapiens* | UniProtKB | P26012 |

**Table 2**

| **Protein Name** | **Organism** | **Database** | **Accession Number** |
|---|---|---|---|
| Paxillin (PXN) | *Homo sapiens* | Alphafold DB | AF-P49023-F1-v4 |
| Vinculin (VCL) | *Homo sapiens* | Alphafold DB | AF-P18206-F1-v4 |
| Talin-1 (TLN1) | *Homo sapiens* | Alphafold DB | AF-Q9Y490-F1-v4 |
| Talin-2 (TLN2) | *Homo sapiens* | Alphafold DB | AF-Q9Y4G6-F1-v4 |

| | | | |
|---|---|---|---|
| Zyxin (ZYX) | *Homo sapiens* | Alphafold DB | AF-Q15942-F1-v4 |
| Focal Adhesion Kinase 1 (FAK1) | *Homo sapiens* | Alphafold DB | AF-Q05397-F1-v4 |
| Focal Adhesion Kinase 2 (FAK2) | *Homo sapiens* | Alphafold DB | AF-Q14289-F1-v4 |
| Proto-oncogene tyrosine-protein kinase SRC (c-SRC) | *Homo sapiens* | Alphafold DB | AF-P41240-F1-v4 |
| Integrin Alpha 1 (ITGA1) | *Homo sapiens* | Alphafold DB | AF-P56199-F1-v4 |
| Integrin Alpha 2 (ITGA2) | *Homo sapiens* | Alphafold DB | AF-P17301-F1-v4 |
| Integrin Alpha 3 (ITGA3) | *Homo sapiens* | Alphafold DB | AF-P26006-F1-v4 |
| Integrin Alpha 4 (ITGA4) | *Homo sapiens* | Alphafold DB | AF-P13612-F1-v4 |
| Integrin Alpha 5 (ITGA5) | *Homo sapiens* | Alphafold DB | AF-P08648-F1-v4 |
| Integrin Alpha 6 (ITGA6) | *Homo sapiens* | Alphafold DB | AF-P23229-F1-v4 |
| Integrin Alpha 7 (ITGA7) | *Homo sapiens* | Alphafold DB | AF-Q13683-F1-v4 |
| Integrin Alpha 8 (ITGA8) | *Homo sapiens* | Alphafold DB | AF-P53708-F1-v4 |
| Integrin Alpha 9 (ITGA9) | *Homo sapiens* | Alphafold DB | AF-Q13797-F1-v4 |
| Integrin Alpha 10 (ITGA10) | *Homo sapiens* | Alphafold DB | AF-O75578-F1-v4 |
| Integrin Alpha 11 (ITGA11) | *Homo sapiens* | Alphafold DB | AF-Q9UKX5-F1-v4 |
| Integrin Alpha D (ITGAD) | *Homo sapiens* | Alphafold DB | AF-Q13349-F1-v4 |
| Integrin Alpha E (ITGAE) | *Homo sapiens* | Alphafold DB | AF-P38570-F1-v4 |
| Integrin Alpha L (ITGAL) | *Homo sapiens* | Alphafold DB | AF-P20701-F1-v4 |
| Integrin Alpha M (ITGAM) | *Homo sapiens* | Alphafold DB | AF-P11215-F1-v4 |
| Integrin Alpha V (ITGAV) | *Homo sapiens* | Alphafold DB | AF-P06756-F1-v4 |
| Integrin Alpha-IIb (ITGA2B) | *Homo sapiens* | Alphafold DB | AF-P08514-F1-v4 |

| | | | |
|---|---|---|---|
| Integrin Alpha X (ITGAX) | *Homo sapiens* | Alphafold DB | AF-P20702-F1-v4 |
| Integrin Beta 1 (ITGB1) | *Homo sapiens* | Alphafold DB | AF-P05556-F1-v4 |
| Integrin Beta 2 (ITGB2) | *Homo sapiens* | Alphafold DB | AF-P05107-F1-v4 |
| Integrin Beta 3 (ITGB3) | *Homo sapiens* | Alphafold DB | AF-P05106-F1-v4 |
| Integrin Beta 4 (ITGB4) | *Homo sapiens* | Alphafold DB | AF-P16144-F1-v4 |
| Integrin Beta 5 (ITGB5) | *Homo sapiens* | Alphafold DB | AF-P18084-F1-v4 |
| Integrin Beta 6 (ITGB6) | *Homo sapiens* | Alphafold DB | AF-P18564-F1-v4 |
| Integrin Beta 7 (ITGB7) | *Homo sapiens* | Alphafold DB | AF-P26010-F1-v4 |
| IIntegrin Beta 8 (ITGB8) | *Homo sapiens* | Alphafold DB | AF-P26012-F1-v4 |

**Table 3**

| Protein Name | Organism | Database | Accession Number |
|---|---|---|---|
| Paxillin (PXN) | *Homo sapiens* | UniProtKB | P49023 |
| Leupaxin (LPXN) | *Homo sapiens* | UniProtKB | O60711 |
| LIM and senescent cell antigen-like containing domain protein 1 (LIMS1) | *Homo sapiens* | UniProtKB | P48059 |
| LIM and senescent cell antigen-like containing domain protein 2 (LIMS2) | *Homo sapiens* | UniProtKB | Q7Z4I7 |
| PDZ and LIM domain protein 7 (PDLIM7) | *Homo sapiens* | UniProtKB | Q9NR12 |
| Testin (TES) | *Homo sapiens* | UniProtKB | Q9UGI8 |
| Zyxin (ZYX) | *Homo sapiens* | UniProtKB | Q15942 |
| Paxillin (PXN) | *Homo sapiens* | Alphafold | AF-P49023-F1-v4 |
| Leupaxin (LPXN) | *Homo sapiens* | Alphafold | AF-O60711-F1-v4 |
| LIM and senescent cell antigen-like containing domain protein 1 (LIMS1) | *Homo sapiens* | Alphafold | AF-P48059-F1-v4 |

| | | | |
|---|---|---|---|
| LIM and senescent cell antigen-like containing domain protein 2 (LIMS2) | *Homo sapiens* | Alphafold | AF-Q7Z4I7-F1-v4 |
| PDZ and LIM domain protein 7 (PDLIM7) | *Homo sapiens* | Alphafold | AF-Q9NR12-F1-v4 |
| Testin (TES) | *Homo sapiens* | Alphafold | AF-Q9UGI8-F1-v4 |
| Zyxin (ZYX) | *Homo sapiens* | Alphafold | AF-Q15942-F1-v4 |

### Supplemental Video Legends

**Video S1: PaxillinB Co-Localizes with Actin at Dynamic Ventral Surface Structures During Cell Migration**
Representative spinning disc confocal timelapse of *paxB⁻ Dictyostelium* cells expressing GFP:PaxillinB and RFP:Lifeact. Left panel shows GFP:PaxillinB, middle panel shows RFP:Lifeact and right panel shows merged movies. Images taken every 5 sec for 10 min; 10 fps.

**Video S2: TIRF Microscopy Shows PaxillinB Punctae Form at the Ventral Surface During Cell Migration**
Representative total internal reflection fluorescence timelapse of *paxB⁻ Dictyostelium* cells expressing GFP:PaxillinB. Images taken every 3 sec for 3 min; 10 fps.

**Video S3: Perturbation of Actin Polymerization Perturbs PaxillinB Punctae Formation**
Representative laser scanning confocal timelapse of *paxB⁻ Dictyostelium* cells expressing GFP:PaxillinB and RFP:Lifeact with LatrunculinA spiked-in to perturb actin polymerization during the movie. Left panel shows GFP:Paxillin, middle panel shows RFP:Lifeact and right panel shows merged movies. Images taken every 5 sec for 10 min; 10 fps.

**Video S4: PaxillinB Co-Localizes with TalinB at Adhesion Structures During Cell Migration**
Representative spinning disc confocal timelapse of *paxB⁻ Dictyostelium* cells expressing GFP:TalinB and mScarlet:PaxillinB. Left panel shows GFP:TalinB, middle panel shows mScarlet:PaxillinB and right panel shows merged movies. Images taken every 5 sec for 10 min; 10 fps.

**Video S5: PaxillinB Co-Localizes with VinculinB at Adhesion Structures During Cell Migration**
Representative spinning disc confocal timelapse of *paxB⁻ Dictyostelium* cells expressing GFP:PaxiilinB and mApple:VinculinB Left panel shows GFP:PaxillinB, middle panel shows mApplet:VinculinB and right panel shows merged movies. Images taken every 3 sec for 3 min; 10 fps.
